# Metabolic control of adult neural stem cell self-renewal by the mitochondrial protease YME1L

**DOI:** 10.1101/2021.08.18.456709

**Authors:** Gulzar A. Wani, Hans-Georg Sprenger, Kristiano Ndoci, Srikanth Chandragiri, Richard James Acton, Désirée Schatton, Sandra M.V. Kochan, Vignesh Sakthivelu, Milica Jevtic, Jens M. Seeger, Stefan Müller, Patrick Giavalisco, Elena I. Rugarli, Elisa Motori, Thomas Langer, Matteo Bergami

**Author notes:** Correspondence to: Matteo Bergami CECAD-University Hospital of Cologne, Germany. Lead contact: Matteo Bergami CECAD-University Hospital of Cologne, Germany.

## Abstract

The transition between quiescence and activation in neural stem and progenitor cells (NSPCs) is coupled to reversible changes in energy metabolism with key implications for life-long NSPC self-renewal and neurogenesis. How this metabolic plasticity is ensured between NSPC activity states is unclear. We found that a state-dependent rewiring of the mitochondrial proteome by the peptidase YME1L is required to preserve NSPC self-renewal in the adult brain. YME1L-mediated proteome rewiring regulates the rate of fatty acid oxidation (FAO) for replenishing Krebs cycle intermediates and dNTP precursors, which are required to sustain NSPC amplification. *Yme1l* deletion irreversibly shifts the metabolic profile of NSPCs away from a FAO-dependent state resulting in defective self-renewal, premature differentiation and NSPC pool depletion. Our results disclose an important role for YME1L in coordinating the switch between metabolic states of NSPCs and suggest that NSPC fate is regulated by compartmentalized changes in protein network dynamics.

## Introduction

Neural stem and progenitor cells (NSPCs) are retained exclusively in few restricted regions of the adult mammalian brain, where they generate new neurons that contribute to specific forms of brain plasticity life-long (Goncalves et al., 2016). Maintenance of this long-lived pool of NSPCs into adulthood is guaranteed by their capability to reversibly switch between proliferative and quiescent states, which confers protection from damage but also prevents irreversible NSPC pool depletion (Navarro Negredo et al., 2020).

The activity state of adult NSPCs is regulated on multiple levels. Cell-autonomously, quality control mechanisms ensuring protein homeostasis like the proteasome and lysosomal-autophagic systems are regulated differentially between quiescent and activated states, and can be manipulated to control NSPC fate (Leeman et al., 2018; Morrow et al., 2020; Schaffner et al., 2018). Likewise, specific metabolic programs including changes in lipid metabolism, reactive oxygen species (ROS) signalling, redox state, glutaminolysis and mitochondrial oxidative phosphorylation (OXPHOS) mark the switch between cellular stages along the embryonic and adult NSPC lineage (Adusumilli et al., 2021; Ahlqvist et al., 2012; Beckervordersandforth et al., 2017; Homem et al., 2014; Khacho et al., 2016; Knobloch et al., 2013; Knobloch et al., 2017; Namba et al., 2020; Prozorovski et al., 2008; Stoll et al., 2015; Xie et al., 2016), suggesting that a rewiring of energy metabolism plays important roles over NSPC fate decisions as it has been proposed for hematopoietic, immune and cancer cells (Mehta et al., 2017; Nakamura-Ishizu et al., 2020; Puleston et al., 2017; Snaebjornsson et al., 2020). Yet, besides overall state-dependent changes in nuclear gene transcription (Llorens-Bobadilla et al., 2015; Shin et al., 2015) it remains unclear how the reversibility of these metabolic programs is coordinated at the level of single organelles in adult NSPCs.

In this study, we simultaneously investigated the changes in energy metabolism and mitochondrial protein network dynamics underlying the activity of adult NSPCs. By using a combination of unbiased omics approaches and conditional mouse models, we identified the *i*-AAA protease YME1L for being central in acutely shaping the mitochondrial proteome of NSPCs between active and quiescent states. Specifically, we demonstrated that YME1L contributes to rewire mitochondrial metabolism to sustain NSPC proliferation. By genetically manipulating YME1L activity *in vitro* and *in vivo*, we showed that this effect is independent from mitochondrial dynamics. Rather, this rewiring process impacts the rate of mitochondrial fatty acid catabolism (FAO) to sustain the production of tricarboxylic acid (TCA) cycle intermediates and nucleotide biosynthesis, which we showed to be necessary for NSPC proliferation. By lineage analysis of NSPC at the single clone level *in vivo*, we demonstrated that lack of YME1L affects stem cell fate by limiting self-renewal and promoting premature differentiation, ultimately causing NSPC pool depletion.

## Results

### YME1L proteolytic activity mirrors opposed metabolic states in adult NSPCs

To reveal dynamics of protein networks underlying reversible, state-dependent changes in fuel utilization of adult NSPCs, we combined an unbiased proteomic approach with metabolomics (Figure 1A). NSPCs were isolated from the adult hippocampal sub-granular zone (SGZ) and maintained *in vitro* in either active proliferation (aNSPC) or quiescence (qNSPC), the latter state induced via addition of bone morphogenetic protein 4 (BMP4) (Mira et al., 2010) (Figure 1A and 1B). By principal component (PCA) and protein distribution analyses, the proteome of qNSPCs could be easily separated from that of aNSPCs and ex-qNSPCs (i.e., qNSPCs in which BMP4 had been withdrawn to disclose reversible changes in the proteome) (Figures S1A and S1B). Furthermore, we validated the expression of well-established markers matching with quiescent and active NSPC states (Figure S1C), as well as the ability of aNSPCs to differentiate into beta-3 tubulin+ neurons following growth factors withdrawal from the proliferation media (Figure S1D). Ingenuity Pathway Analysis (IPA) of this proteomic dataset revealed a number of differentially regulated proteins with pathways associated to cell cycle control, nucleotide excision repair, gene transcription and purine biosynthesis that were preferentially enriched in aNSPCs (Figure 1C). In contrast, besides expected categories linked to phagosome/lysosome and autophagy (Leeman et al., 2018), qNSPCs appeared particularly enriched in metabolic pathways, specifically mitochondrial OXPHOS as well as fatty acid beta-oxidation (FAO) (Figure 1C), a catabolic pathway which has been recently implicated in regulating adult NSPC behaviour (Knobloch et al., 2017; Stoll et al., 2015). Hierarchical clustering of our dataset revealed conspicuous yet largely reversible changes in the mitochondrial proteome (i.e., proteins annotated according to MitoCarta 3.0) (Rath et al., 2021) of NSPCs shifting from proliferation to quiescence within virtually each mitochondrial compartment (Figure 1D). We identified 307 mitochondrial proteins for being significantly changed (adjusted p-value < 0.05) between aNSPCs and qNSPCs, of which 64.2% were up-regulated and 35.8% down-regulated. In particular, the steady-state levels of most FAO enzymes, several OXPHOS proteins as well as TCA cycle enzymes appeared differentially regulated between aNSPCs and qNSPCs (Figure 1D). While assessment of mitochondrial membrane potential between qNSPCs and aNSPCs disclosed overall unchanged levels (Figure S1E), single protein analysis confirmed that the relative abundance of many OXPHOS subunits, particularly of complexes I, IV and V as well as that of enzymes regulating mitochondrial FAO and TCA cycle metabolism was significantly higher in qNSPCs (Figures S1F-S1H), while their expression to large degree reversed following BMP4 withdrawal. Consistent with these data, metabolomics tracing analysis following feeding of qNSPCs with either ^13^C_6_-Glucose or ^13^C_16_-Palmitate confirmed a higher flux for fatty acids as a carbon source to fuel TCA cycle metabolism in comparison to glucose (Figure S1I).

**Figure 1.**
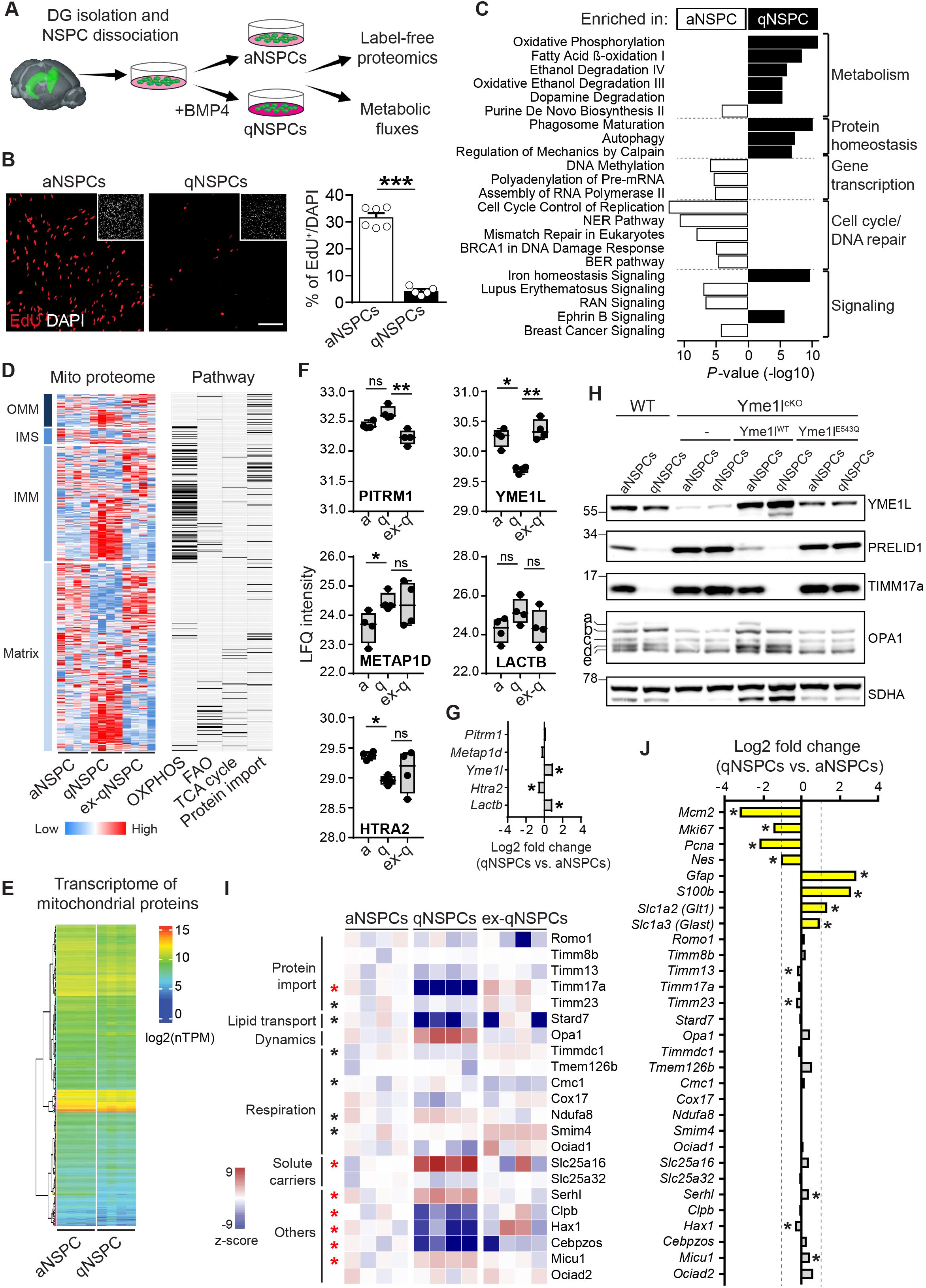
NSPC state-dependent regulation of YME1L activity. **(A)** Experimental setting used for isolating NSPCs from the adult DG and for their maintenance as either actively proliferating (aNSPCs) or quiescent cells (qNSPCs) following BMP4 addition. **(B)** EdU labelling of aNSPCs and qNSPCs and relative quantification (n= 6 and 5 experiments; unpaired t-test). Bar, 80 µm. **(C)** IPA pathways most significantly enriched in either aNSPCs (10 most significant) or qNSPCs (10 most significant). Categories selected for having a p value ≤ 0.01 and a fold enrichment of at least 0.5. **(D)** Cluster analysis of the mitochondrial proteome in aNSPCs, qNSPCs and ex-qNSPCs, displayed according to mitochondrial compartment (OMM, outer mitochondrial membrane; IMS, intermembrane space; IMM, inner mitochondrial membrane). Normalized expression levels are depicted as heat-maps for each quantified protein and scaled row-wise (n=4 experiments per condition). The right panel shows the matched distribution of the quantified proteins and enzymes associated to the indicated pathways. **(E)** Cluster analysis of the transcriptome corresponding to mitochondrial proteins in aNSPCs and qNSPCs (n=4 experiments per condition; nTPM, normalized transcripts per million). **(F)** Abundance levels (label-free quantification, LFQ intensities) of the 5 mitochondrial proteases displaying significant state-dependent changes in NSPCs (n=4 experiments per condition; non-parametric Kruskal-Wallis test). **(G)** Normalized changes in mRNA expression levels between qNSPCs and aNSPCs of the 5 mito-proteases shown in F. **(H)** Immunoblot of wild-type and Yme1l^cKO^ NSPCs with or without expression of Yme1l^WT^ or Yme1l^E543Q^, maintained as active or quiescent (data representative from two independent experiments). **(I)** Z-score heat-maps of normalized LFQ intensities of detected class I putative YME1L substrates in aNSPCs, qNSPCs and ex-qNSPCs, categorized according to their function. Significant changes (n=4 experiments per condition; FDR-adjusted ≤ 0.05) are indicated with an asterisk (red asterisk: qNSPCs significant vs both aNSPCs and ex-NSPCs; black asterisk: qNSPCs significant vs only one other category). **(J)** Fold change of mRNA levels of the indicated genes between qNSPCs and aNSPCs (n=4 experiments; FDR-adjusted ≤ 0.05). Means ± SEM; *, *P* < 0.05; **, *P* < 0.01; ***, *P* < 0.005; ns, not significant. See also Figures S1 and S2.

To understand whether these proteome dynamics may mirror corresponding changes in gene expression, we performed a comparative transcriptomic analysis of active versus quiescent NSPCs (Figures 1E and S1J). Analysis of genes encoding for mitochondrial proteins revealed that a total of 241 genes underwent significant changes (i.e., with a p-value < 0.05 and a *log*2 (fold change) > 0.5), with 39.4% being up-regulated and 60.6% down-regulated. Interestingly, the observed proteomic changes of several TCA cycle enzymes and OXPHOS subunits, and in particular of most FAO proteins, were poorly mirrored at the transcript level (Figure S1F-S1H and S1K). In contrast, analysis of glycolytic enzymes - which reside in the cytosol - disclosed a much higher correlation between protein and corresponding mRNA levels (Figure S1K). This suggests that acute switches in NSPC metabolic states may be regulated by compartmentalized changes in mitochondrial protein networks independent from gene transcription.

The turn-over of several classes of proteins in mitochondria is regulated by the proteolytic activity of proteases distributed across sub-mitochondrial compartments (Quiros et al., 2015). While mitochondria utilize these proteases to broadly preserve mitochondrial proteostasis and function, accruing evidence supports additional roles in acutely shaping the mitochondrial proteome to match specific metabolic needs (Deshwal et al., 2020). Among all detected mitochondrial proteases in our proteomic dataset, only five of them (i.e., PITRM1, YME1L, METAP1D, LACTB and HTRA2) disclosed significant changes between the examined NSPC activity states (Figures 1F and S2A). Of these, a striking state-dependent and fully reversible switch in the levels of the *i*-AAA peptidase YME1L caught our attention, with qNSPCs displaying a significant reduction of this protein as compared to both aNSPCs and ex-qNSPCs (Figure 1F). We independently validated YME1L protein to accumulate at decreased levels in qNSPCs (Figure S2B), however this change was not mirrored by a transcriptional down-regulation, as *Yme1l* mRNA appeared mildly yet significantly increased during quiescence (Figure 1G). YME1L is an inner mitochondrial membrane peptidase whose mutation in humans causes brain disorders, intellectual disability and optic nerve atrophy (Hartmann et al., 2016). YME1L has been implicated in the proteolytic control of numerous mitochondrial substrates (MacVicar et al., 2019) including the GTPase OPA1, whose regulated processing by both YME1L and OMA1 controls inner mitochondrial fusion dynamics (Anand et al., 2014; Griparic et al., 2007; Song et al., 2007). Because of its proteolytic activity, YME1L has been proposed to link mitochondrial dynamics to proteostasis regulation in somatic tissues (Mishra et al., 2014; Sprenger et al., 2019; Wai et al., 2015) and recently to metabolic reprogramming of mitochondria during hypoxia and starvation in cell lines (MacVicar et al., 2019). YME1L mode-of-action involves its own autocatalytic processing when highly active, which leads to a conspicuous reduction of both, YME1L direct substrates and YME1L protein itself (Hartmann et al., 2016; MacVicar et al., 2019). Intriguingly, previously validated mitochondrial targets of YME1L (e.g., PRELID1 and TIMM17a) accumulated at strongly reduced levels in qNSPCs (Figure S2B), thus suggesting increased YME1L-mediated proteolytic degradation of substrate proteins in qNSPCs alongside increased YME1L autocatalytic turnover. To validate this possibility, we took advantage of *Yme1l*^lox/lox^ mice (Anand et al., 2014) to isolate and grow *in vitro* NSPCs followed by treatment with an AAV expressing Cre-GFP to induce *Yme1l* gene deletion (hereafter referred to as Yme1l^cKO^ NSPCs) (Figure S2C). As expected, Cre expression resulted in the virtual disappearance of the endogenous YME1L protein alongside an increase in the steady-state levels of its substrates PRELID1 and TIMM17a, regardless of NSPC growing conditions (i.e., cells maintained in either proliferating or quiescent media) (Figure 1H). Importantly, while AAV-mediated over-expression of wild-type YME1L (Yme1l^WT^) in Yme1l^cKO^ NSPCs restored PRELID1 and TIMM17a proteolytic processing in a state-dependent manner (that is, higher processing in qNSPCs), expression of a mutated, proteolytically-inactive YME1L variant (Yme1l^E543Q^) (MacVicar et al., 2019), proved ineffective (Figure 1H). Thus, we conclude that a differential YME1L proteolytic activity in adult NSPCs reflects the acquisition of distinct metabolic states.

### YME1L is required for mitochondrial proteome rewiring between NSPC states

We next examined in our proteomic dataset the levels of additional mitochondrial proteins which have been recently proposed as likely substrates of YME1L in mouse embryonic fibroblasts (MEFs) (MacVicar et al., 2019). Of these, we identified 22 recently annotated class I targets (namely, putative substrates fulfilling stringent criteria) and 24 additional putative substrates previously designated as class II (MacVicar et al., 2019). These include enzymes regulating mitochondrial protein import, lipid transport, dynamics (i.e., the GTPase OPA1), solute carriers, respiration and other metabolic functions (Figures 1I and S2D). Several of these proteins, particularly those designated as class I, accumulated at visibly reduced levels specifically in qNSPCs (Figure 1I). However, transcriptomic analysis revealed that collectively there were minor or no changes in the corresponding mRNA levels of these genes between qNSPCs and aNSPCs (Figure 1J). In contrast, non-mitochondrial markers known to be preferentially expressed in either active or quiescent states displayed matched changes at both mRNA and protein levels (Figure 1J and S1C). Interestingly, some of the putative YME1L substrates appeared to be only moderately reduced or in certain cases (particularly in class II) even upregulated (Figures 1I and S2D). This may reflect a certain cell type-specificity of the proteolytic activity of YME1L in adult NSPCs as compared to MEFs (MacVicar et al., 2019), or further layers of regulation beyond protein turnover as exemplified by several of class II putative substrates, whose expression in NSPCs consistently differed at both the mRNA and protein levels (Figures S2D and S2E).

To gain further insights into the potential cell-type specificity of identified class I YME1L substrates in NSPCs, we took advantage of Yme1l^cKO^ NSPCs and performed a proteomic analysis utilizing as controls *Yme1l*^lox/lox^ NSPCs treated with an AAV expressing only GFP (Figure S2C). At the mitochondrial level, *Yme1l* deletion caused the significant accumulation of most previously annotated class I substrates which we identified for being specifically downregulated in wild-type qNSPCs (Figure 2A), consistent with their turn-over being under the proteolytic control of YME1L. We defined this subset as NSPC-specific, putative YME1L substrates (Figure S2F). Accordingly, transcriptomic analysis of Yme1l^cKO^ NSPCs revealed that the mRNA levels of these substrates remained virtually unchanged (Figure 2B). By contrast, very few of the previously annotated class II substrates underwent any visible accumulation at the protein level following *Yme1l* deletion (Figure S2G), confirming that YME1L substrate specificity may indeed be regulated in a cell type-specific manner.

**Figure 2.**
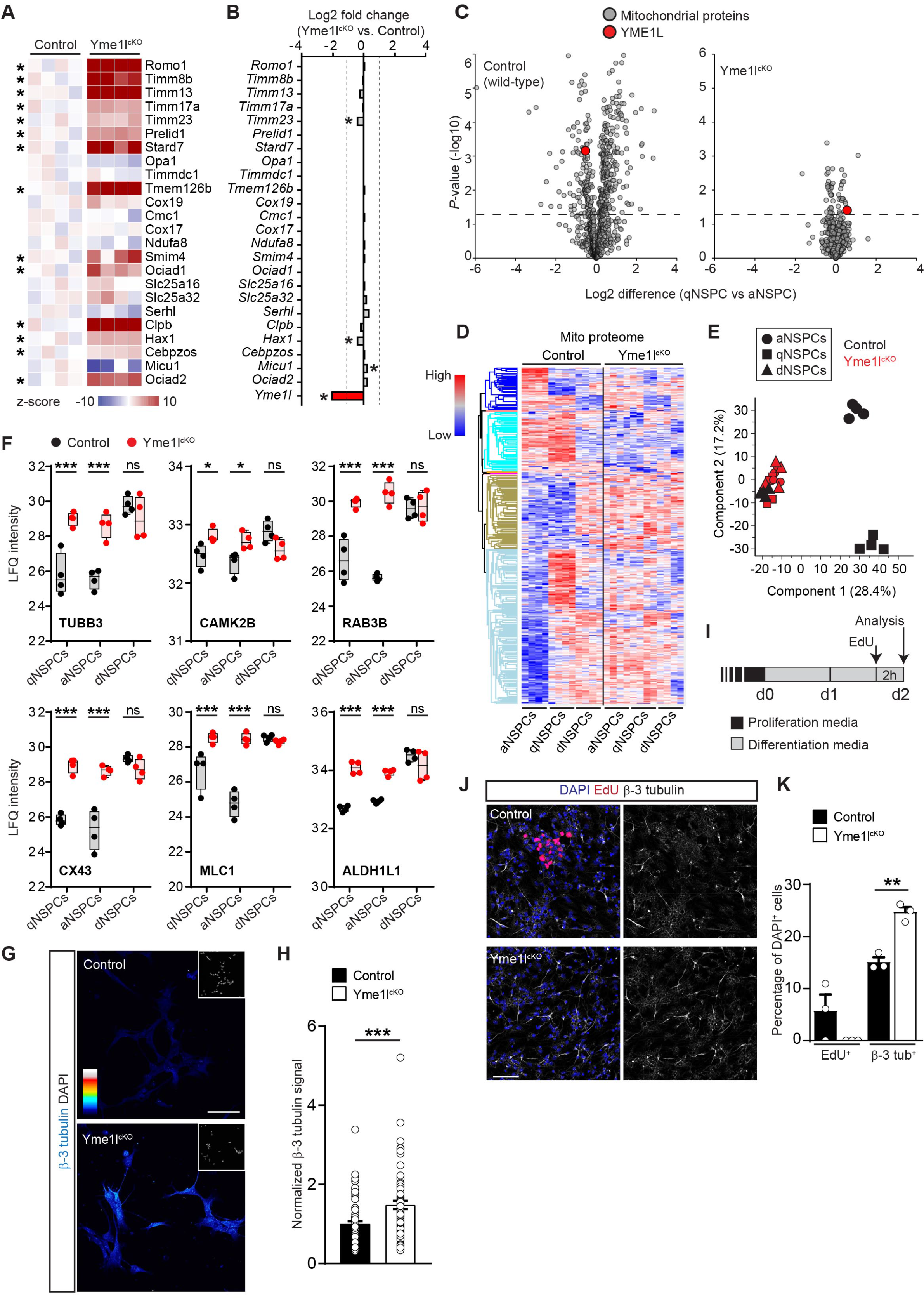
YME1L-dependent rewiring of the mitochondrial proteome in adult NSPCs. **(A)** Heat maps of normalized LFQ intensities of detected class I YME1L putative substrates in control and Yme1l^cKO^ NSPCs. Significant changes are indicated with an asterisk (n= 4 experiments per condition; FDR-adjusted ≤ 0.05). **(B)** mRNA levels of class I YME1L putative substrates in Yme1l^cKO^ versus control NSPCs (n= 3 experiments; FDR-adjusted ≤ 0.05). **(C)** Volcano plots showing the changes in the mitochondrial proteome of quiescent vs active states in control (wild-type) and Yme1l^cKO^ NSPCs. Cut-off line set at –log10 (*P*-value) = 1.3 (n= 4 experiments per dataset). **(D)** Whole-proteome cluster analysis of control and Yme1l^cKO^ NSPCs maintained under active, quiescent and differentiating conditions. Normalized expression levels are depicted as heat-maps for each quantified protein and scaled row-wise (n= 4 experiments per condition). **(E)** PCA plot of control and Yme1l^cKO^ aNSPCs, qNSPCs and dNSPCs proteomic datasets (n= 4 experiments per condition). **(F)** Abundance levels (LFQ intensities) of selected neuronal (TUBB3, CAMK2A and MAP2) and astrocytic (CX43, GS and ALDH1L1) markers in control and Yme1l^cKO^ aNSPCs, qNSPCs and dNSPCs (n= 4 experiments per condition; non-parametric Kruskal-Wallis test). **(G)** Representative examples of control and Yme1l^cKO^ NSPCs maintained under proliferative media and immunostained for the neuronal marker β-3 tubulin (TUBB3). Images are presented in pseudocolors. Insets show location of nuclei. Bar, 50 μm. **(H)** Quantification of β-3 tubulin immunoreactivity in control and Yme1l^cKO^ NSPCs maintained under proliferative media (n= 66 and 69 cells pooled from 2 independent experiments; Welch’s t-test). **(I)** Experimental timeline used to induce short NSPC differentiation combined with EdU labeling for the experiment shown in J. **(J)** Representative examples of control and Yme1l^cKO^ NSPCs exposed for 2 days to differentiation media and immunostained for the EdU and for the neuronal marker β-3 tubulin. Bar, 80 μm. **(K)** Quantification of the experiment shown in J (n= 3 independent experiments; Welch’s t-test). Means ± SEM; *, *P* < 0.05; **, *P* < 0.01; ***, *P* < 0.005; ns, not significant. See also Figure S2.

Next, we investigated the significance of the NSPC state-dependent asymmetry in YME1L activity. We reasoned that *Yme1l* conditional deletion would provide a valid approach to irreversibly interfere with the pronounced YME1L proteolytic activity that specifically identifies the quiescent state in NSPCs (Figure 1H). *Yme1l* deletion was validated at the protein level and by the marked accumulation of one of its direct targets (PRELID1) (Figures S2H and S2I). As expected, we also observed stress-induced OPA1 processing by the peptidase OMA1 as indicated by the accumulation of S-OPA1 forms c and e and the virtual lack of S-OPA1 form d, which results from specific YME1L cleavage (Figure S2H) (Anand et al., 2014). In line with an increased OPA1 processing, morphological analysis of Yme1l^cKO^ NSPCs *in vitro* revealed a fragmented and condensed mitochondrial network in comparison to control NSPCs (Figure S2J), which was also mirrored by a visibly reduced proliferative capacity in Yme1l^cKO^ NSPCs (Figures S2K and S2L). Conspicuously, NSPC proliferation was not restored in double-knockout Yme1l/Oma1^cKO^ NSPCs (Figures S2K and S2L), in which OPA1 cleavage (by both YME1L and OMA1) is virtually abolished and thus mitochondrial fusion as well as network tubulation restored (Figure S2J) (Anand et al., 2014; Wai et al., 2015). Thus, YME1L appears to be required for sustaining adult NSPC proliferation *in vitro* via mechanisms seemingly independent of its role in OPA1 processing and the regulation of mitochondrial morphology (Iwata et al., 2020; Khacho et al., 2016).

Analysis of the proteomic landscape in Yme1l^cKO^ NSPCs revealed the unexpected lack of any significant shift in the mitochondrial proteome between active (proliferating) and quiescent conditions, as otherwise observed in wild-type NSPCs (Figure 2C). Intriguingly, hierarchical cluster analysis and PCA showed that, regardless of culture media conditions, *Yme1l* deletion led to a broad rewiring of the mitochondrial proteome beyond class I substrates, which was consistent with the acquisition of a mitochondrial “state” more similar to control NSPCs shortly exposed (2 days) to pro-differentiating conditions (dNSPCs) (Figure 2D and 2E). In marked contrast to Yme1l^cKO^ NSPCs, deletion of the protease *Oma1* only led to negligible changes in the mitochondrial proteome of NSPCs (Figure S2M), indicating that the observed proteomic rewiring was specific to *Yme1l* deletion. This apparent shift towards a differentiated-like state of Yme1l^cKO^ NSPCs (Figure 2E) was reflected by the up-regulation of some neuron- (TUBB3, CAMK2B and RAB3B) as well as astrocyte-specific (CX43, MLC1, ALDH1L1) markers even when Yme1l^cKO^ NSPCs were specifically maintained in proliferating or quiescent media, contrary to control NSPCs in which this pattern was exclusively induced after beginning of the differentiation protocol (Figure 2F). Although Yme1l^cKO^ NSPCs did not spontaneously differentiate into neurons (as assessed by morphology and marker expression), analysis of TUBB3 (i.e., β-3 tubulin) immunoreactivity in cells maintained under proliferative media revealed a consistent up-regulation in comparison to control NSPCs (Figure 2G and 2H), suggesting that *Yme1l* deletion may facilitate neuronal differentiation under proper media conditions. Supporting this prediction, Yme1l^cKO^ NSPCs exposed shortly (2 days) to differentiation media (Figure 2I) underwent an accelerated differentiation into β-3 tubulin^+^ neurons at the expenses of residual proliferation (Figures 2J and 2K). Together, these data indicate that *Yme1l* deletion is sufficient to elicit broad changes in mitochondrial proteome dynamics consistent with the reduction in NSPC proliferation and a shift towards a differentiated-like state.

### Loss of YME1L in NSPCs impairs fatty acid-dependent metabolic flux into the TCA cycle and leads to dNTP pool depletion

Direct comparison of the mitochondrial proteome between Yme1l^cKO^ and control NSPCs (maintained under either quiescent or active conditions) disclosed a down-regulation of proteins falling into the FAO category (Figure 3A). In contrast, deletion of *Oma1* did not visibly affect the expression levels of FAO enzymes (Figure S3A). Likewise, analysis of the mitochondrial proteome in NSPCs lacking *Mitofusin1* (*Mfn1*) or *Mfn2* revealed that a collective down-regulation of FAO proteins alongside defects in NSPC proliferation did not necessarily reflect a cellular state characterized by mitochondrial fragmentation (Figures S3B-3D). Specifically, Yme1l^cKO^ NSPCs failed in up-regulating FAO proteins when exposed to quiescent conditions (Figure 3B), despite unchanged or even slightly up-regulated mRNA levels of the corresponding FAO genes (Figure 3C). Thus, while FAO enzymes do not appear to be under the direct control of YME1L proteolytic activity, these data suggest that *Yme1l* deletion prevents the proteomic rewiring underlying key switches in mitochondrial fuel utilization between NSPC activity states. To validate this hypothesis, we first utilized SeaHorse analysis to examine oxygen consumption rates in Yme1l^cKO^ NSPCs. While basal respiration was not affected as compared to control NSPCs, we found a selective impairment in maximal and spared respiratory capacities when cells were fed with palmitate, but not glucose (Figures 3D and 3E). Interestingly, glucose feeding experiments showed that Yme1l^cKO^ NSPCs could achieve even higher peaks of respiration rates than control NSPCs (Figures 3E) and that there were no deficits in the NAD/NADH ratio (Figure S3E), indicating that OXPHOS capacity in these cells was likely not primarily impaired. Supporting this notion, mitochondrial membrane potential assessed by Tetramethylrhodamine methyl ester (TMRM) revealed no major changes in the average signal intensity between Yme1l^cKO^ and control NSPCs, despite obvious alterations in mitochondrial morphology (Figures 3F and 3G). Likewise, ultrastructural analysis of Yme1l^cKO^ NSPCs disclosed that fragmented mitochondria retained cristae (Figure 3H) indicating that, similar to other cellular systems (Sprenger et al., 2019; Wai et al., 2015), *Yme1l* deletion is not associated to mitochondrial dysfunction and aberrant ultrastructure in adult NSPCs.

**Figure 3.**
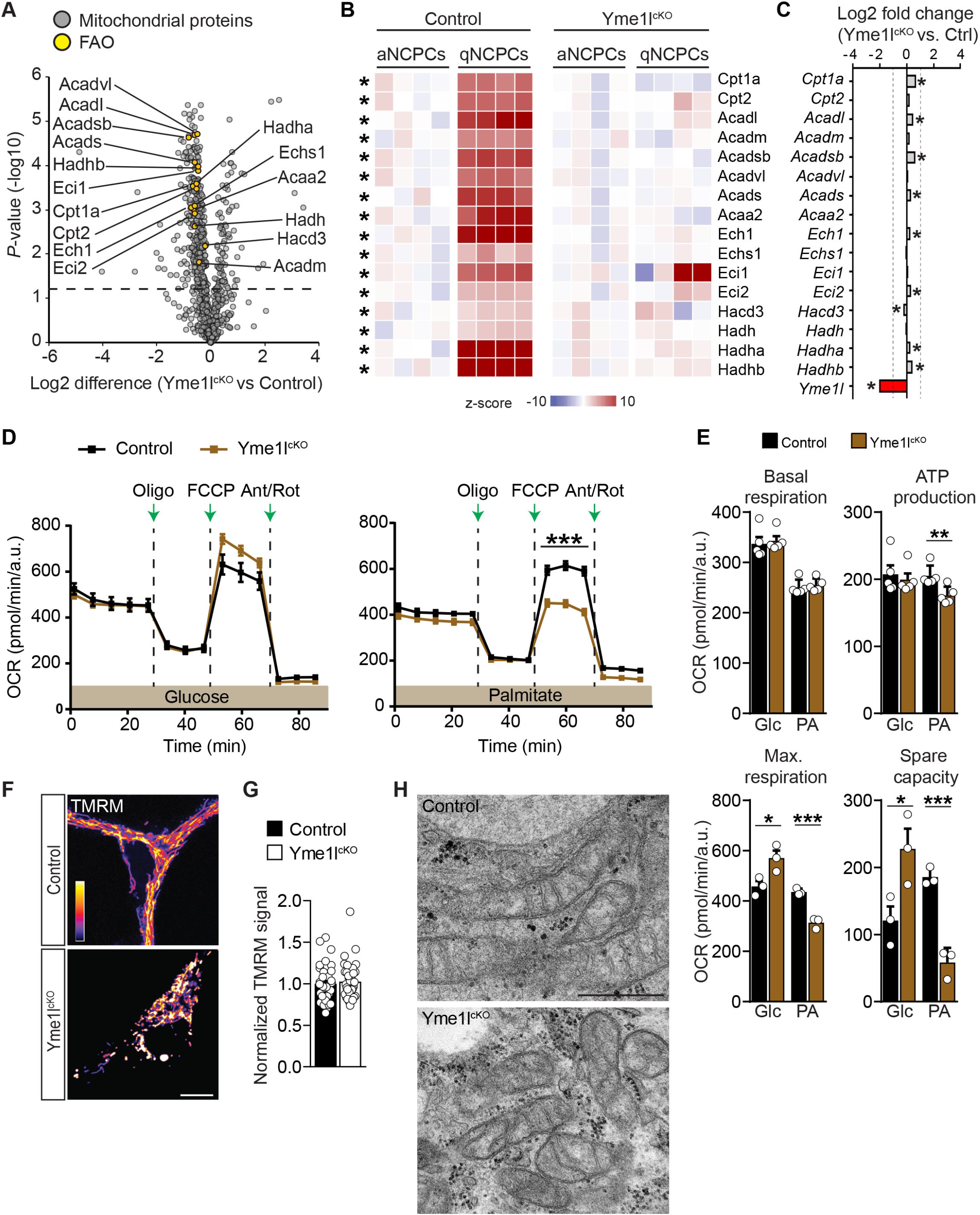
Impaired FAO in adult NSPCs lacking YME1L. **(A)** Volcano plot showing the changes of mitochondrial FAO-associated proteins in Yme1l^cKO^ NSPCs (n= 4 experiments). Cut-off line set at –log10 (*P*-value) = 1.3. **(B)** Heat-maps of normalized LFQ intensities of FAO-associated proteins in control and Yme1l^cKO^ NSPCs maintained in either proliferative or quiescent conditions (n= 4 experiments per condition). **(C)** mRNA levels of FAO enzymes in Yme1l^cKO^ versus control NSPCs (n= 3 experiments; FDR-adjusted ≤ 0.05). **(D)** Oxygen consumption rates of control and Yme1l^cKO^ NSPCs fed with either glucose or palmitate (n= 30 repetitions pooled from 3 independent experiments; Holm-Sidak multiple t-test). **(E)** SeaHorse analysis of basal respiration, ATP production, maximal respiration and spare respiratory capacity in Yme1l^cKO^ NSPCs fed with either glucose or palmitate (n= 3 to 5 independent experiments; unpaired t-test). **(F)** Examples of NSPCs of the indicated genotype following incubation with Tetramethylrhodamine methyl ester (TMRM, 25 nM) for 10 minutes. Bar, 10 µm. **(G)** Quantification of TMRM signal intensity in control and Yme1l^cKO^ NSPCs (n= 34-36 cells pooled from 2 independent experiments; Welch’s t-test). **(H)** Electron micrographs of mitochondria in control and Yme1l^cKO^ NSPCs. Bar, 500 nm. Means ± SEM; *, *P* < 0.05; **, *P* < 0.01; ***, *P* < 0.005; ns, not significant. See also Figure S3.

To further dissect the consequences of *Yme1l* deletion for NSPC mitochondrial metabolism, we analysed isotope labelling of TCA cycle intermediates and associated newly-synthetized amino acids by metabolomics following feeding with either ^13^C_16_-Palmitate or ^13^C_6_-Glucose. We found that labelling of most TCA cycle metabolites was consistently and specifically reduced in Yme1l^cKO^ NSPCs fed with ^13^C_16_-Palmitate (Figure 4A), while of the quantified amino acids only aspartate and glutamate appeared to be significantly affected (Figure 4B). In contrast, the ^13^C_6_-Glucose flux into the TCA cycle of Yme1l^cKO^ NSPCs appeared to remain stable or even somewhat enhanced (Figures 4A and 4B). Importantly, quantification of total TCA cycle metabolites revealed no collective changes besides a reduction in the content of citrate and isocitrate (Figure S3I), arguing against a general dysfunction of the TCA cycle and confirming an overall lower flux of specifically ^13^C_16_-Palmitate into TCA cycle metabolites (Figures 4A). Also, besides mild changes in total glycolytic metabolites (Figure S3H) Yme1l^cKO^ NSPCs fed with glucose displayed a higher labelling of alanine, pyruvate and lactate (Figure S3F), which was reflected by a higher extracellular acidification rate (ECAR) when cells were examined by Seahorse (Figures S3G). Together, these data point to a general metabolic rewiring taking place in Yme1l^cKO^ NSPCs, and support a defective feeding into the TCA cycle of specifically fatty acid carbon units.

**Figure 4.**
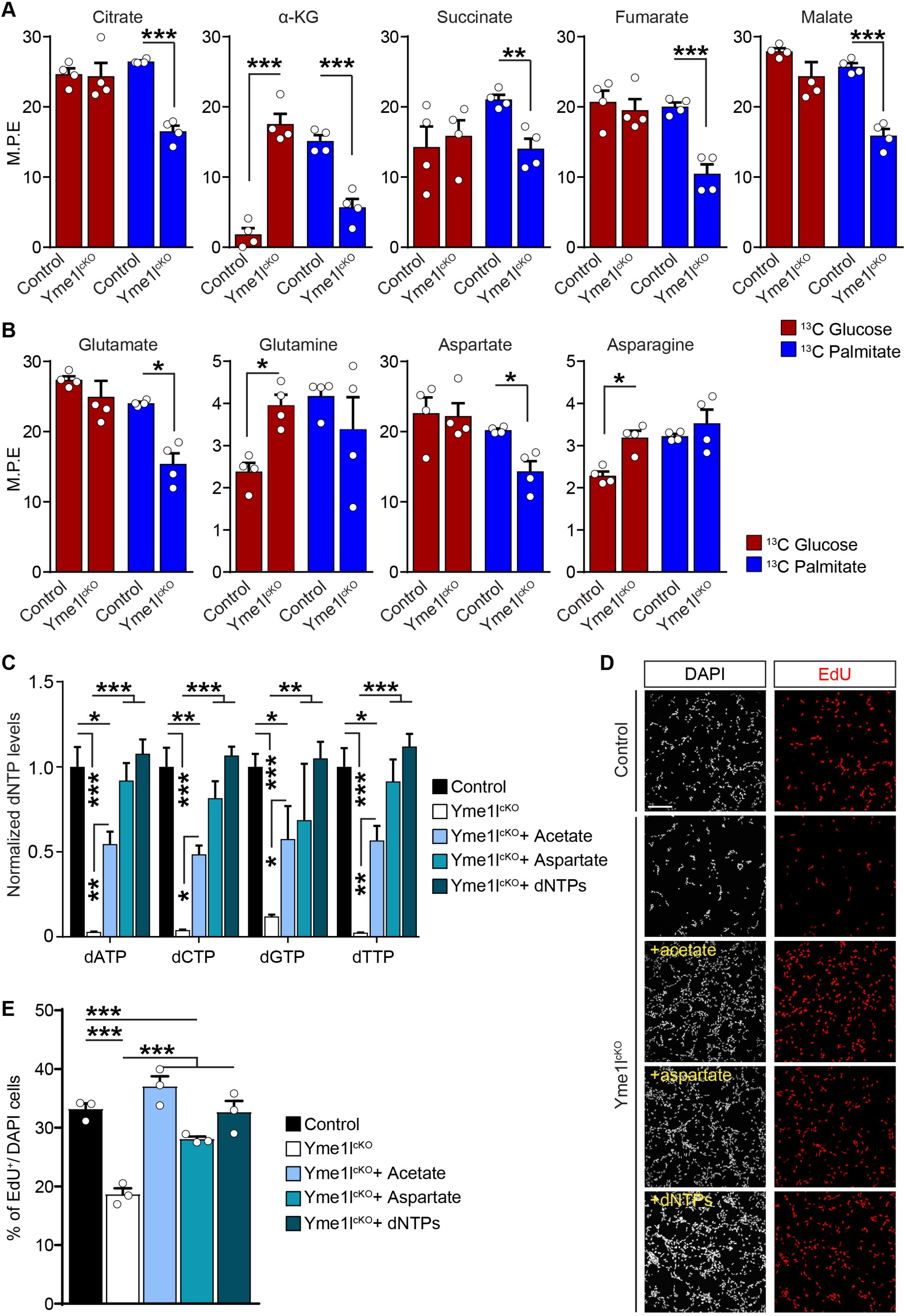
Loss of YME1L in NSPCs impairs fatty-acid carbon feeding into the TCA cycle resulting in dNTP pool depletion. **(A)** Quantification of TCA cycle metabolites following either U-^13^C_6_-Glucose or U-^13^C_6_-Palmitate isotope labelling in control and Yme1l^cKO^ NSPCs (n= 4 experiments per condition; Holm-Sidak multiple t-test). M.P.E., Molar Percent Enrichment (calculated for each isotope). **(B)** Quantification of newly-synthetized amino acids following either U-^13^C_6_-Glucose or U-^13^C_6_-Palmitate isotope labelling in control and Yme1l^cKO^ NSPCs (n= 4 experiments per condition; Holm-Sidak multiple t-test). **(C)** Quantification of dNTP levels in control and Yme1l^cKO^ NSPCs with or without supplementation with the indicated compounds (n= n= 4 experiments per condition; Two-way Anova followed by Tukey’s multiple comparison test). **(D)** Examples showing EdU incorporation in Yme1l^cKO^ NSPC cultures with or without supplementation of exogenous dNTPs, acetate or aspartate. Bar, 80 µm. **(E)** Quantification of NSPC proliferation with or without supplementation of exogenous dNTPs, acetate or aspartate (n= n= 3 experiments per condition; One-way Anova followed by Holm-Sidak’s correction). Means ± SEM; *, *P* < 0.05; **, *P* < 0.01; ***, *P* < 0.005; ns, not significant. See also Figure S3.

The selective reduction in aspartate and glutamate in NSPCs that were fed with palmitate (Figure 4B) raised the possibility that the proliferation defects observed in Yme1l^cKO^ NSPCs may, at least in part, result from reduced levels of nucleotides, which are required for cell proliferation and specifically rely on the precursors aspartate and glutamate for their biosynthesis (Schoors et al., 2015). Indeed, the steady-state levels of purine and pyrimidine deoxyribonucleotides (dNTPs) were markedly reduced in absence of YME1L (Figure 4C). Consistent with a defective FAO-dependent supply of carbon units for dNTPs biosynthesis, supplementation of exogenous dNTPs, acetate or even aspartate to Yme1l^cKO^ NSPCs mostly restored dNTP levels, albeit at different degrees (Figure 4C). To assess whether manipulation of dNTP content would be sufficient to reactivate NSPC proliferation following *Yme1l* deletion, we then examined EdU incorporation while maintaining NSPCs in proliferating media. Intriguingly, the tested compounds significantly improved Yme1l^cKO^ NSPC proliferative capacity (Figures 4D and 4E) indicating that, *in vitro*, manipulation of the dNTP pool can effectively compensate for the metabolic alterations of Yme1l^cKO^ NSPC. Thus, *Yme1l* deletion drives NSPCs away from FAO-dependent metabolic states, causing the subsequent depletion of dNTP precursors required to sustain NSPC proliferation.

### YME1L is required for adult NSPC proliferation *in vivo*

To circumvent the limitations imposed by an *in vitro* system and defined media conditions, we next addressed the role of YME1L in adult NSPCs *in vivo* by crossing *Yme1l*^lox/lox^ mice with a line harbouring an hGFAP-driven, tamoxifen-inducible Cre^ER^ recombinase (Chow et al., 2008). To reveal putative changes in NSPC mitochondrial morphology induced by Cre recombination, which would be indicative of *Yme1l* deletion, we bred the resulting line with a mitochondrial-targeted yellow fluorescent protein (mtYFP) reporter mouse (Sterky et al., 2011) (Figures 5A and S4A). Analysis of NSPCs in the SGZ of the resulting Yme1l^cKO^ mice at 1 month following tamoxifen administration disclosed a striking fragmentation of their mitochondrial network, in contrast to control NSPCs, which retained a heterogeneous yet overall tubular network (Figure 5B). Intriguingly, simultaneous deletion of both *Yme1l* and *Oma1* in double-floxed animals mostly restored a tubular mitochondrial morphology (Figure 5B), validating our results *in vitro* (Figure S2J) that the fragmentation phenotype observed in Yme1l^cKO^ NSPCs is to large degree mediated by stress-induced activation of OMA1 followed by excessive OPA1 processing (Anand et al., 2014; Sprenger et al., 2019; Wai et al., 2015). Next, we examined the proliferative capacity of adult Yme1l^cKO^ NSPCs. By 4 weeks after Cre-mediated recombination, EdU incorporation experiments (Figure 5A) revealed a significant reduction in the number of proliferating NSPCs of Yme1l^cKO^ mice both within the SGZ (Figures 5C and 5D) and sub-ventricular zone (SVZ) lining the lateral ventricles (Figures S4E and S4F), the second major neurogenic niche in the adult murine brain. This finding was corroborated by expression analysis of the endogenous proliferative marker Ki67 in SOX2+ NSPCs within the SGZ of the dentate gyrus (Figures S4G-S4I). Interestingly, while no proliferation changes were observed in Oma1^cKO^ mice (Figures S4B-S4D), simultaneous deletion of both *Yme1l* and *Oma1* produced the same defect observed in Yme1l^cKO^ mice (Figures 5C, 5D and S4E-S4I), confirming this phenotype to be mediated by an YME1L-specific metabolic function and ruling out possible primary effects caused solely by alterations in mitochondrial fusion dynamics (Iwata et al., 2020; Khacho et al., 2016). Supporting these findings, AAV-mediated re-expression of wild-type YME1L (Yme1l^WT^) in Yme1l^cKO^ NSPCs *in vitro* (Figures S5A) or *in vivo* via intracranial delivery into the DG (Figures 5E and S5B-S5D) significantly restored NSPC proliferation and mitochondrial morphology, in contrast to expression of a mutated, proteolytically-inactive YME1L variant (Yme1l^E543Q^), which proved ineffective. Interestingly, forced expression of Yme1l^E543Q^ alone was sufficient to reduce NSPC proliferation in control mice (Figure 5E), suggestive of a dominant negative effect. Subsequent analysis of Ki67^+^ cells in Yme1l^cKO^ animals treated with EdU 4 days earlier (Figure S5E) revealed a marked reduction in the percentage of EdU-retaining NSPCs that were still proliferating at the time of sacrifice compared to control mice (12.6% in Yme1l^cKO^ mice vs. 42.9% in controls) (Figures S5F-S5H), demonstrating that lack of YME1L severely affected NSPC pool amplification. To understand whether NSPC proliferation dynamics were already altered at times earlier than 4 weeks following Cre-mediated *Yme1l* deletion, mice were treated with EdU at 5-6 days after tamoxifen administration and examined 4 days later (Figure 5F). Analysis revealed unchanged numbers of Nestin^+^ cells and a trend towards a reduced total density of EdU^+^ cells, which manifested as a significant drop of EdU incorporation into Nestin^+^ cells (Figure 5G and 5H). Thus, these data are consistent with an early proliferative defect in Nestin^+^ NSPC that persisted later on.

**Figure 5.**
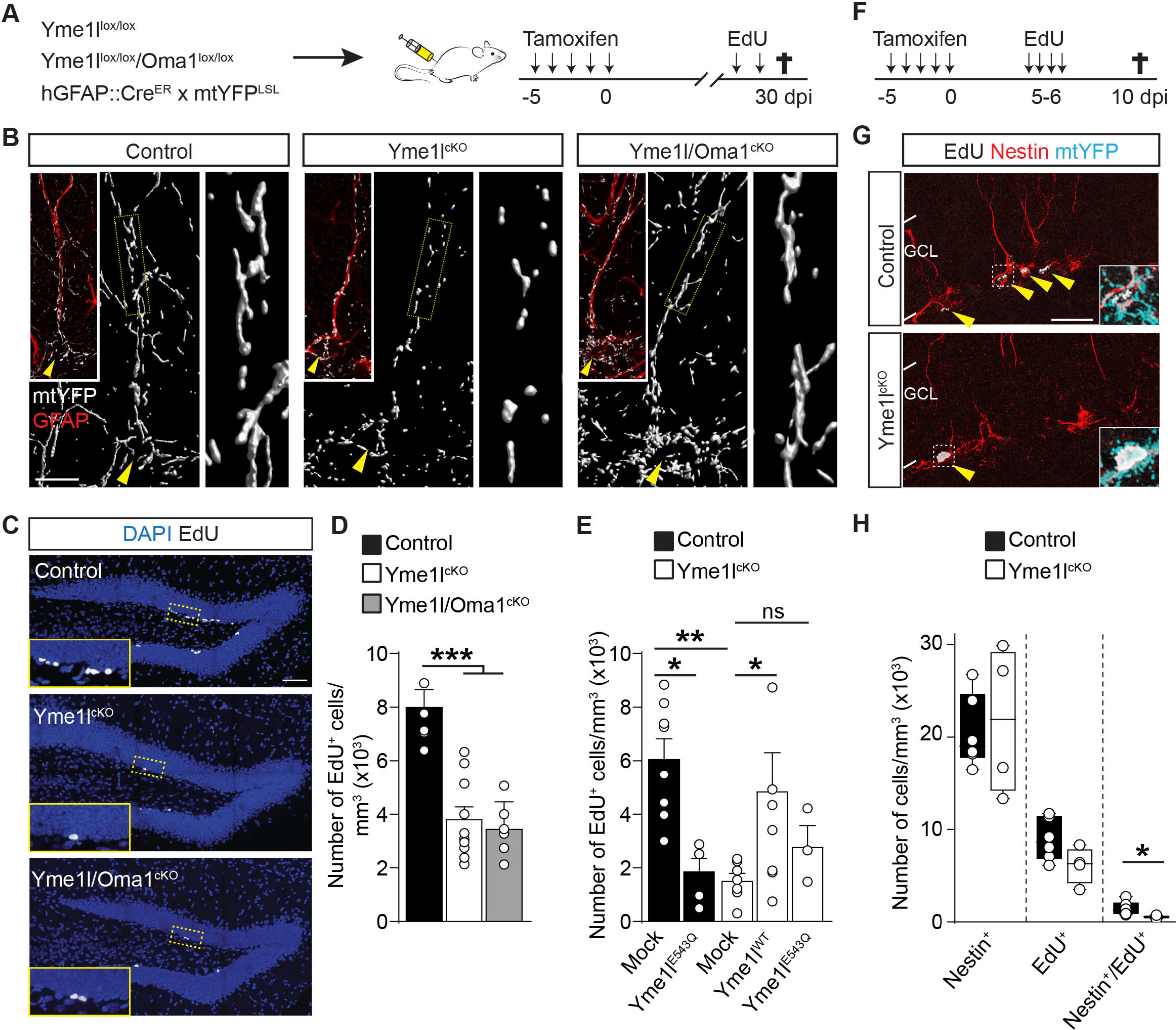
Defective NSPC proliferation in the adult DG of Yme1l^cKO^ mice. **(A)** Experimental design illustrating the tamoxifen-induced conditional deletion of *Yme1l*, or *Yme1l* and *Oma1*, in hGFAP::Cre^ER^ x mtYFP^LSL^ mice. **(B)** Examples of radial glia-like mtYFP^+^/GFAP^+^ NSPCs (see inset) in the DG of the indicated genotypes, showing the morphology of individual mitochondria. Arrowheads points to the cell soma. Right panels show zooms of the boxed areas along the main radial process. Bar, 15 µm. **(C)** Examples of EdU labelling in the DG of the indicated genotypes. Insets show zooms of the boxed areas. Bar, 70 µm. **(D)** Quantification of NSPC proliferation for the indicated genotypes (n= 5-10 mice per group; One-way Anova followed by Holm-Sidak’s multiple comparison test). **(E)** Quantification of NSPC proliferation for the indicated genotypes and conditions following delivery of mock, Yme1l^E543Q^ or Yme1l^WT^ AAVs (n= 3-8 mice per group; One-way Anova followed by Holm-Sidak’s multiple comparison test). **(F)** Experimental design utilized to assess NSPC proliferation at 10 days after tamoxifen-induced recombination in Yme1l^cKO^ mice. **(G)** Examples of Nestin, EdU and mtYFP labelling in the DG of the indicated genotypes. Arrowheads point to EdU^+^ cells. Insets show zooms of individual Nestin^+^/EdU^+^ cells. Bar, 30 µm. **(H)** Quantification of Nestin^+^, EdU^+^ and Nestin^+^/EdU^+^ NSPCs in control and Yme1l^cKO^ mice (n= 4-6 mice per group; Welch’s t-test). Means ± SEM; *, *P* < 0.05; **, *P* < 0.01; ***, *P* < 0.005; ns, not significant. See also Figures S4 and S5.

### *Yme1l* conditional deletion impairs adult NSPC self-renewal and promotes pool depletion

To gain further insights into the consequences of *Yme1l* deletion for NSPC fate, we generated inducible Yme1l^cKO^ mice in which Cre recombination is conditionally controlled by the Nestin promoter (Nestin::Cre^ERT2^) (Lagace et al., 2007) and utilized a cytosolic tdTomato reporter line (Madisen et al., 2010) to fate map any resulting progeny from recombined radial glia-like (RGL) NSPCs. Mice were treated with tamoxifen for 5 consecutive days, in order to label the majority of NSPCs and their immediate neuronal progeny during their first month of maturation (Figure 6A). In line with our EdU experiments (Figure 5D), analysis revealed a drastic reduction in the total density of tdTomato^+^ cells in Yme1l^cKO^ mice as compared to control littermates (Figure 6B). Specifically, the density of Sox2^+^ RGL NSPCs (namely, type I cells) as well as that of Tbr2^+^ (type II) cells and newly-generated Dcx^+^ immature neurons was significantly reduced (Figure 6C, S6A and S6B), confirming that neurogenesis was impaired. In contrast, the density of new neurons transiting from immature to mature stages (Dcx^+^/NeuN^+^) as well as that of NeuN^+^ neurons that had just matured appeared similar between the two groups at this relatively early time of 1 month post-recombination (Figure 6C). Also, the relative proportion of NeuN+ neurons among all neuronal maturational stages was not visibly different compared to that of control mice (Figure 6D), suggesting that neuronal maturation was not primarily affected in absence of *Yme1l* and consistent with a putative progressive depletion of NSPCs in favour of premature differentiation. Supporting this notion, cell density analysis performed at 3 months after tamoxifen administration disclosed a marked impairment in the generation and subsequent network addition of further NeuN^+^ new neurons (Figure 6C), indicating cumulative defects in neurogenesis driven by exhaustion of the NSPC pool.

**Figure 6.**
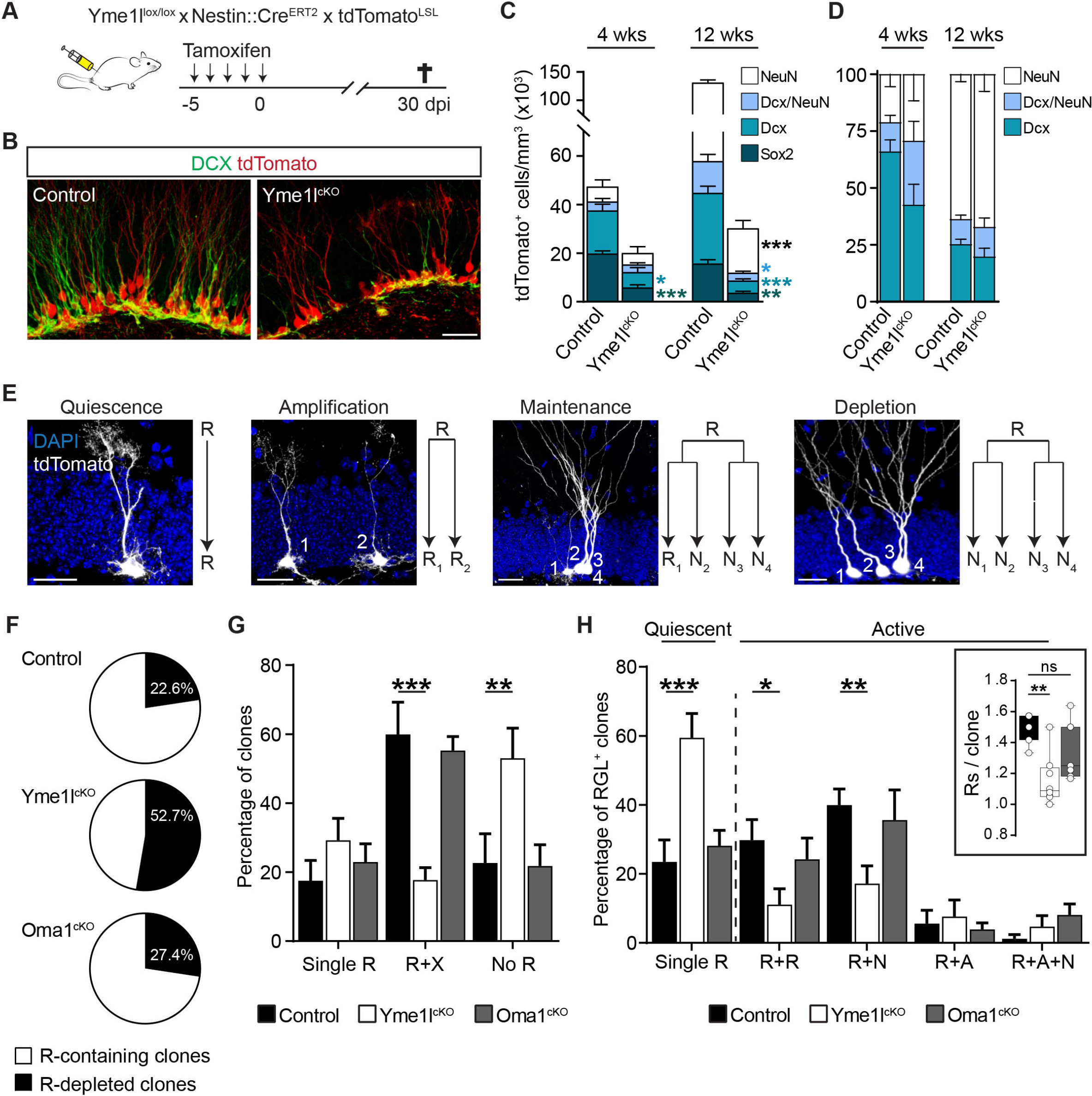
*Yme1l* conditional deletion promotes NSPC pool depletion at the expenses of self-renewal. **(A)** Experimental design illustrating the tamoxifen-induced conditional deletion of *Yme1l* in Yme1l^lox/lox^ x Nestin::Cre^ERT2^ x tdTomato^LSL^ mice. **(B)** Examples showing the amount of tdTomato^+^ cells and Dcx^+^ new neurons by 1 month after tamoxifen administration. Bar, 30 µm. **(C)** Quantification of tdTomato^+^ cells according to their identity (SOX2^+^ RGL NSPCs; Dcx^+^ immature neurons; Dcx^+^/NeuN^+^ maturing neurons; and NeuN^+^ mature neurons) at 4 and 12 weeks after tamoxifen-mediated recombination (n= 3-4 mice per group; Holm-Sidak multiple t-test). **(D)** Percentage of tdTomato^+^ immature and mature neurons at 4 and 12 weeks post-recombination quantified according to C (n= 3-4 mice per group; Holm-Sidak multiple t-test). **(E)** Examples of individual clones at 1 month after tamoxifen administration. The putative lineage outcome of each initially recombined radial glia-like NSPC (R) is shown. Bars, 25 µm. **(F)** Fraction of clones containing (with R) or depleted of (without R) radial glia-like NSPCs for the indicated genotypes (n= 7-8 mice per group). **(G)** Quantification of clones containing single R (quiescent), R+X (self-renewing) and no R (depleted) for the indicated genotypes (n= 7-8 mice per group, Two-way Anova followed by Tukey’s multiple comparison test). **(H)** Proportion of R-containing clones classified according to their cellular composition (n= 7-8 mice per group, Two-way Anova followed by Tukey’s multiple comparison test) (R, radial glia-like NSPC; N, neuron; A, astrocyte). Inset reports on the number of Rs per clone (n= 7-8 animals per group). Means ± SEM; *, *P* < 0.05; **, *P* < 0.01; ***, *P* < 0.005; ns, not significant. See also Figure S6.

To ascertain the specific consequences caused by *Yme1l* deletion for NSPC behaviour, we performed a clonal analysis *in vivo* by targeting few individual RGL NSPCs within each hippocampi (Bonaguidi et al., 2011). Low titre tamoxifen treatment in Yme1l^cKO^ mice resulted in sparse labelling of isolated RGL NSPCs (Figures S6C and S6D), which allowed us to monitor their activity and progression along the lineage at the level of individual clones (Figure 6E). Specifically, cell composition within isolated clones revealed whether recombined RGL NSPCs had remained quiescent, underwent division (symmetrically or asymmetrically) and if they became eventually depleted from the clone over the course 1 month (Figure 6D). Detailed analysis of clone composition showed that while control and Oma1^cKO^ tdTomato^+^ clones were in average larger in size (Figure S6E) and possessed a mixed combination of neurons, astrocytes and RGL NSPCs (referred to as R), which is indicative of NSPC being able to self-renew (Figures 6F and 6H), the majority (over 50%) of Yme1l^cKO^ clones were entirely devoid of any R (indicative of NSPC pool depletion) (Figures 6F and 6G). By further examining the remaining fraction of clones still containing Rs, we found that in average Yme1l^cKO^ clones contained 1.15 ± 0.06 Rs per clone (109 Rs in 99 clones in total from 8 animals), in contrast to control (71 Rs in 47 clones in total from 7 animals) and Oma1^cKO^ (117 Rs in 80 clones in total from 8 animals) clones, which contained in average 1.5 ± 0.03 and 1.35 ± 0.07 Rs per clone, respectively (Figure 6H, inset). In particular, the relative proportion of R doublets (i.e., purely amplifying clones) as well as neurogenic clones (i.e., containing R and neurons) were significantly reduced in Yme1l^cKO^ animals (11% amplifying clones in Yme1l^cKO^ versus 30% in control mice; and 22% neurogenic clones in Yme1l^cKO^ versus ∼41% in control mice) (Figure 6H). Thus, *Yme1l* deletion compromises adult NSPC self-renewal capacity and pool maintenance, ultimately impacting neurogenesis.

## Discussion

Owing to their functional heterogeneity, the fate of NSPCs in the mouse DG can have different outcomes. In young adult mice, a significant fraction of NSPCs undergoes a limited number of self-renewing divisions interspersed by temporary quiescent states, before being eventually depleted (Bonaguidi et al., 2011; Encinas et al., 2011; Pilz et al., 2018). However, populations of NSPCs with extended periods of quiescence favouring long-term self-renewing capacity exist, and likely critically contribute to continuous neurogenesis during adulthood (Bottes et al., 2021; Harris et al., 2021; Ibrayeva et al., 2021). By examining changes in the mitochondrial proteome mirroring the acquisition of active and quiescent states, we have identified an unexpected role for the protease YME1L in maintaining the self-renewing potential of adult NSPCs seemingly independent from its role in balancing mitochondrial dynamics via OPA1 processing (Anand et al., 2014; Wai et al., 2015). Mitochondrial fission/fusion dynamics have an instructing role in regulating the fate of NSPCs and early post-mitotic cells during embryonic cortical development, with fusion promoting self-renewal and mitochondrial network fragmentation favouring differentiation (Iwata et al., 2020; Khacho et al., 2016). Intriguingly, by simultaneously manipulating *Yme1l* and *Oma1* in adult NSPCs we restored a tubular mitochondrial network in absence of YME1L and revealed YME1L-specific effects beyond changes in mitochondrial morphology. These are consistent with a broader proteolytic function of YME1L in shaping the mitochondrial proteome to instruct and accommodate specific metabolic adaptations of NSPCs, as recently observed in cell lines exposed to hypoxia or starvation, conditions that promote a metabolic shift towards OXPHOS-independent, glycolytic growth and are of relevance for certain types of cancer (MacVicar et al., 2019). Different from immortalized embryonic fibroblasts and cancer cell lines, however, we found a significant degree of NSPC specificity in the proteolytic activity of YME1L, as revealed by the assessment of putative direct substrates accumulating following *Yme1l* deletion. For instance, most previously annotated class II substrates (MacVicar et al., 2019) did not appear to be under the direct (or unique) control of YME1L processing in adult NSPCs. These include the mitochondrial membrane transporter SLC25A33, whose up-regulation in *Yme1l*-deficient cell lines was recently linked to mitochondrial DNA release into the cytosol with ensuing inflammatory response and marked activation of interferon-stimulated genes (Sprenger et al., 2021), of which we found no evidence in adult Yme1l^cKO^ NSPCs (Figure S3J). Thus, although it remains conceivable that additional NSPC-specific substrates may exist beyond those initially proposed by MacVicar et al., it appears that YME1L activity is differentially regulated in a cell type- and, possibly, stimulus-specific manner.

In our system, proteomics and metabolomics showed that YME1L is required to preserve NSPC activity states, as lack of YME1L caused the irreversible shift of the mitochondrial proteome towards a FAO-independent, differentiated-like state. Accordingly, Yme1l^cKO^ NSPCs displayed a drop in proliferation *in vitro* and *in vivo*. *In vitro*, the resulting reduction in FAO rates observed in absence of YME1L was found to be responsible for the overall drop in proliferation, even when NSPCs were exposed to growth factors, indicating that Yme1l^cKO^ NSPCs attained an altered metabolic state unable to respond to proliferative stimuli. Mechanistically, we identified the feeding of fatty acid carbon units into TCA cycle-derived intermediates that were required to sustain the dNTP pool as the bottleneck causing this proliferative phenotype, as exogenous replenishment of either dNTPs or their precursors was sufficient to revert proliferation. Besides impairing proliferation, *Yme1l* deletion in NSPCs resulted in an accelerated propensity to differentiate into neurons when growth factors were withdrawn from the growing media *in vitro* and once RGL NSPCs became activated *in vivo*. Strikingly, deletion of *Yme1l* at the single clone level *in vivo* led to a phenotype reminiscent of the genetic ablation of FAO, which drives NSPC exit from quiescence, potentiating terminal neurogenic symmetric divisions at the expenses of self-renewal (Knobloch et al., 2017; Xie et al., 2016). Likewise, we found that over 50% of examined Yme1l^cKO^ clones contained neurons (and astrocytes) but were devoid of any RGL NSPC, which is consistent with NSPC pool depletion and in line with the metabolic alterations induced by defective FAO.

Together, our results reveal YME1L for playing a critical role in acutely shaping the mitochondrial proteome of NSPCs, adding an important layer of regulation in the mechanisms governing NSPC metabolic state transitions beyond potential changes in gene expression (Beckervordersandforth et al., 2017; Llorens-Bobadilla et al., 2015; Shin et al., 2015). Importantly, our data indicate that switches in fuel utilization of adult NSPCs can be coordinated by compartmentalized dynamics in protein networks, and suggest that the activity of mitochondrial proteases critically contributes to regulate this form of metabolic plasticity.

### Limitations of study

While our data emphasize the role of mitochondrial proteome rewiring in regulating NSPC activity, an obvious limitation of our study is that the proteomics and metabolomics data presented here were exclusively obtained from adult NSPCs isolated and maintained *in vitro*. Owing to technological limitations that currently prevent to efficiently discriminate and sort distinct NSPC activity states (e.g., active and quiescent) from brain tissue, it thus remains unclear to which extent NSPCs adjust their mitochondrial proteome and metabolome in a state-dependent manner *in vivo*. Assessing these changes *in vivo* may pose further challenges on account of distinct depths of quiescence acquired by NSPCs during adulthood (Bottes et al., 2021; Harris et al., 2021; Ibrayeva et al., 2021). Yet, with the advent of refined strategies suitable for single-cell mass spectrometry-based proteomics (Brunner et al., 2021), further investigations should aim at reconstructing the proteomic landscape heterogeneity of NSPC activity states *in vivo* as it has been shown at the transcriptomic level utilizing single-cell RNA-seq approaches (Bottes et al., 2021; Harris et al., 2021; Llorens-Bobadilla et al., 2015; Shin et al., 2015).

## Author contributions

Conceptualization, G.A.W. and M.B.; Methodology, G.A.W., R.J.A, J.M.S., S.M., P.G., E.M. and M.B.; Investigation, G.A.W., H-G.S., K.N., S.C., D.S. and M.B.; Formal analysis, G.A.W., H-G.S., K.N., R.J.A, S.M.V.K., D.S., S.M., P.G., E.M. and M.B.; Writing -Original Draft, M.B.; Writing – Review and Editing, G.A.W., H-G.S., D.S., P.G., E.I.R., E.M., T.L. and M.B.; Resources, V.S., M.J., T.L. and M.B.; Funding Acquisition, E.I.R., E.M., T.L. and M.B.; Supervision, M.B.; Project Administration, M.B.

## Acknowledgments

We thank N.G. Larsson for providing mitoYFP floxed-stop mice. B. Fernando and T. Öztürk for excellent technical assistance. J. Matutat, G. Piper, D. Schneider and the other members of the CECAD in vivo facility for excellent assistance. A. Schauss and all members of the CECAD imaging facility for assistance with microscopes. C. Frese and the team of the CECAD proteomics core facility for assistance. J. Altmüller and the team of the Cologne Center for Genomics (CCG) for assistance with RNA-Seq procedures. This work was supported by the Deutsche Forschungsgemeinschaft (SFB1218 - Grant No. 269925409, SFB1451 - Grant No. 431549029 and CECAD EXC 2030 – Grant No. 390661388) and European Research Council (ERC-StG-2015, grant number 67844) to M.B.; the Deutsche Forschungsgemeinschaft (SFB1218 - Grant No. 269925409) to T.L. and E.I.R.; and by the German-Israeli-Project (DIP, RA1028/10-1) and the Max-Planck-Society to T.L. E.M. was supported by an Advanced Postdoc Grant (Deutsche Forschungsgemeinschaft, SFB1218 - Grant No. 269925409).

## Author Information

The authors declare no competing financial interests.

## STAR methods

### Lead contact and materials availability

Further information and requests for resources and reagents should be directed to and will be fulfilled by the Lead Contact, Matteo Bergami. All unique/stable reagents generated in this study are available from the Lead Contact without restrictions. There are restrictions to the availability of mice due to MTA.

### Experimental model and subject details

Six to 8-week old C57BL/6 and transgenic mice of mixed genders were used in this study. Mice were housed in groups of up to 5 animals per cage supplied with standard pellet food and water *ad libitum* with a 12 h light/dark cycle, while temperature was controlled to 21-22°C. Mice carrying the loxP-flanked genes *Yme1l*^fl/fl^, *Oma1*^fl/fl^ or both (Anand et al., 2014) were crossed with the inducible hGFAP-Cre^ERTM^ (Chow et al., 2008) line and subsequently to the Cre-dependent mitochondrial-targeted mtYFP reporter (Sterky et al., 2011). For clonal analysis experiments, *Yme1l*^fl/fl^ or *Oma1*^fl/fl^ mice were crossed with the Nestin-Cre^ERT2^ line (Lagace et al., 2007) in combination with the inducible tdTomato reporter (Madisen et al., 2010). Mice carrying the loxP-flanked genes *Mfn1*^fl/fl^ and *Mfn2*^fl/fl^ (Lee et al., 2012) were utilized exclusively for preparation of NSPC cultures maintained *in vitro*. All experimental procedures were performed in agreement with the European Union and German guidelines and were approved by the State Government of North Rhine Westphalia.

### Method details

#### Tamoxifen and EdU treatments

Mice were intraperitoneally injected with 4-hydroxytamoxifen (40 mg/ml dissolved in 90% corn oil and 10% ethanol) once a day for a 5 consecutive days. For clonal analysis, mice received a single injection of 0.4 mg. The exact time frames of individual experiments are indicated in the text and figures. To examine NSPC proliferation, mice were given 5-ethynyl-2-deoxyuridine (EdU) via i.p. injections (25mg/ml, stock solution dissolved in 0.9% saline) and sacrificed 2 hours or 4 days after the last injection.

#### Stereotactic procedures and viral injections

Mice were anesthetized by intraperitoneal injection of a ketamine/xylazine mixture (100 mg/kg body weight ketamine, 10 mg/kg body weight xylazine), treated subcutaneously with Carprofen (5 mg/kg) and fixed in a stereotactic frame provided with a heating pad. A portion of the skull covering the somatosensory cortex (from Bregma: caudal: -2.0; lateral: 1.5) was thinned with a dental drill avoiding to disturb the underlying vasculature and small craniotomy sufficient to allow penetration of a glass capillary performed. For virus injection a finely pulled glass capillary was then inserted through the dura (-1.9 to -1.8 from Bregma) and a total of about 500 nl of virus were slowly infused via a manual syringe (Narishige) in multiple vertical steps spaced by 50 µm each during a time window of 10-20 minutes. After infusion, the capillary was left in place for few additional minutes to allow complete diffusion of the virus. After capillary removal, the scalp was sutured and mice were placed on a warm heating pad until full recovery. Physical conditions of the animals were monitored daily to improve their welfare before euthanize them.

#### Viral production

Helper-free AAV vectors were either obtained from Addgene or produced according to standard manufacturer’s instructions (Cell Biolabs) as previously described (Göbel et al., 2020). Briefly, 293AAV cells were transiently transfected with a transfer plasmid carrying the desired transgenes along with a packing plasmid encoding the AAV1 capsid proteins and a helper plasmid, using the calcium phosphate method. Crude viral supernatants were obtained via lysing cells in PBS by freeze-thaw cycles in a dry ice/ethanol bath. The AAV1 vectors were purified by discontinuous iodixanol gradient ultracentrifugation (24h at 32,000 rpm and 4°C) and concentrated using Amicon ultra-15 centrifugal filter unites. Genomic titres were determined by real-time qPCR. For generation of AAVs expressing Yme1l1^WT^ and Yme1l1^E543Q^, the corresponding previously described sequences (MacVicar et al., 2019) were subcloned into the destination AAV plasmid and sequenced for validation before producing the final AAV.

#### Immunohistochemistry

Mice were anesthetized by intraperitoneal injection of a ketamine/xylazine mixture (130 mg/kg body weight ketamine, 10 mg/kg body weight xylazine), transcardially perfused with 4% PFA in PBS and the brain isolated. Following overnight post-fixation, coronal brain sections (50 µm thick) were prepared using a vibratome (Leica, VT1000 S) and permeabilized in 1% Triton X-100 in PBS for 10 min at RT, followed by brief incubation in 5% BSA and 0.3% Triton X-100 in PBS before overnight immunodetection with primary antibodies diluted in blocking buffer at 4°C on an orbital shaker. The next day, sections were rinsed in PBS 3x 10 min and incubated for 2h at RT with the respective fluorophore-conjugated secondary antibodies diluted in 3% BSA. After washing and nuclear counterstaining with 4’,6-diamidino-2-phenylindole (DAPI, ThermoFisher, 3 µM), sections were mounted on microscopic slides using Aqua Poly/Mount (Polysciences). The following primary antibodies were used: chicken anti-GFP (1:500, Aves Labs, GFP-1020), rabbit anti-RFP (1:500, Rockland, 600401379), rabbit anti-GFAP (1:500, Millipore, ab5804), mouse anti-GFAP (1:500, Millipore, MAB360), mouse anti-Tbr2 (1:300, Abcam, ab23345), guineapig anti-Dcx (1:1000, Millipore, AB2253), mouse anti-NeuN (1:300, Millipore, MAB377) and mouse anti-Nestin (1:500, Millipore, MAB353). The following secondary antibodies were used (raised in donkey): Alexa Fluor 488-, Alexa Fluor 546-, Alexa Fluor 647-conjugated secondary antibodies to rabbit, mouse, chicken and rat (1:1000, Jackson ImmunoResearch).

#### Image analysis and quantification

All images were acquired utilizing a SP8 Confocal microscope (Leica Microsystems) equipped with 20x (NA 0.75), 40x (NA 1.3), 63x (NA 1.4) and 100x (NA 1.3) immersion objectives, white light laser and multiple HyD detectors. For quantification of cell proliferation and neurogenesis *in vivo*, six to eight coronal brain sections per brain corresponding to similar anatomical locations across mice were used. All acquired z-stack images (LAS software) were converted into TIFF files and analysis was performed off-line using the ImageJ software (National Institute of Health, Bethesda, United States). Cells counting was performed manually by using the Cell Counter plug in and by normalizing the number of marker+ cells over the volume of the DG or SVZ (measured area multiplied by the inter-stack interval). The total number of RGL in the DG was obtained by examining their unique morphological features, which makes them well distinguishable from other cell types (short radial process spanning the granule cell layer, small body in the SGZ), and positivity for the marker GFAP. Likewise, the number of neurons was assessed by unique morphological features (round cell body and a visible dendritic arbor) as well as by the positivity for immature (Dcx) or more mature markers (NeuN). For clonal analysis, the whole hippocampus was investigated through serial brain sections (thickness 75µm), which were first screened for TdTomato+ cells with a 20x oil objective and then z-stack acquisition of regions containing positive cells taken with a 20x and a 63x oil objectives. For quantification, regions including the molecular layer (ML), granular cell layer (GCL), sub granular zone (SGZ) and Hilus bordering the SGZ of the DG were considered. A radius of 150 μm from the RGL cell within the clone was used to determine the spatial limits of the clone itself (Bonaguidi et al., 2011). Clones were categorized according to the presence or absence of RGL and by the composition of other cell types (neurons and astrocytes). After initial validation by marker expression of possible TdTomato+ cell types (see above), cell identity in individual clones was determined based on morphology: RGL with a triangular-shaped soma located in the SGZ, a short radial process branching in the inner molecular layer and absence of any axon; neurons for having round cell body located in in the GCL and clearly visible apical dendritic processes; and protoplasmic astrocytes, for having a bushy morphology irrespective of their locations. For imaging of mitochondria in RGL *in vivo*, mice bearing the mtYFP reporter were utilized, and z-stack acquisitions of individual RGL in the upper blade of the DG (identified by location of their soma in the SGZ and the presence of a distinct GFAP+ radial process) taken with a 63x oil objective utilizing an inter-stack interval of 0.3 μm. Following acquisition, images were processed by deconvolution utilizing the Huygens Pro software (Scientific Volume Imaging) and rendered in volumetric views to appreciate changes in mitochondrial morphology.

#### Mitochondrial membrane potential measurements

For membrane potential experiments *in vitro*, seeded NSPCs were incubated with TMRM (25 nM) for 10 minutes in imaging media (124 mM NaCl, 3 mM KCl, 2 mM CaCl2, 1 mM MgCl2, 10 mM HEPES, 10 mM D-glucose and pH 7.4), followed by 1x washing and live imaging utilizing a confocal microscope (inverted Leica SP8, 40x oil objective) and a 560 nm excitation wavelength with a laser power inferior to 1%. Cells were randomly selected for their positivity to the AAV-encoded GFP reporter (cytosolic GFP for control cells, nuclear Cre-GFP for knock-out cells) as well as presence of TMRM staining and acquired with identical parameters (laser power, zoom, resolution, scanning speed, pinhole, digital gain and offset) for all conditions. Total TMRM signal was then quantified via ImageJ for individual cells and normalized to the covered TMRM area to compensate for differences in mitochondrial network morphology and size.

#### Electron microscopy

Cells were grown on aclar foil and fixed with a pre-warmed solution of 2% glutaraldehyde, 2.5% sucrose, 3 mM CaCl2, 100 Mm HEPES, pH 7.4 at RT for 30min and 4°C for other 30 min. Cells were washed with 0.1 M sodium cacodylate buffer (Applichem), incubated with 1%OsO_4_ (Science Services),1.25% sucrose and 10mg/ml potassium ferrocyanid in 0.1M cacodylate buffer for 1 hour on ice, and washed three times with 0.1M cacodylate buffer. Subsequently, cells were dehydrated using ascending ethanol series (50, 70, 90, 100%) for 7 min each at 4°C. Afterwards, cells were incubated with ascending EPON series (Sigma-Aldrich) at 4°C, transferred to fresh EPON for 2h at RT and finally embedded for 72 hours at 62°C. Ultrathin sections of 70 nm were cut using an ultramicrotome (Leica Microsystems, EM-UC7) and a diamond knife (Diatome, Biel, Switzerland). Sections were put on copper grids (mesh 100, Science Service) with formvar film and stained with uranyl acetate for 15 min at 37°C and lead nitrate solution for 4 min. Electron micrographs were taken with a JEM-2100 Plus Transmission Electron Microscope (JEOL), a OneView 4K 16 bit camera (Gatan) and the software DigitalMicrograph (Gatan).

#### NSPCs isolation and cell culture experiments

Primary adult mouse NSPCs were isolated and cultured as previously described (Walker and Kempermann, 2014). In brief, young adult mice (5-6 week old) were quickly sacrificed by cervical dislocation and the brain isolated under sterile conditions. The regions corresponding to the DG or SVZ of the lateral ventricles were microdissected and collected in cold Neurobasal-A. The pooled tissue from 5-6 mice was chopped with a scalpel into small fragments and incubated in pre-warmed Papain (2.5 U/ml), Dispase (1 U/ml) and DNase (250 U/ml) cocktail (PDD) enzyme mix for 20 min at 37°C. The cells were then triturated 10-15 times with a fire-polished Pasteur pipette, collected by centrifugation for 5 min (200g) at room temperature and enriched for the NSPC fraction using a 22% Percoll gradient. Cells were washed twice in Neurobasal-A and plated in proliferation medium containing Neurobasal-A, B27, Glutamax and FGF (1:2500) and EGF (1:2500) into Poly-D-lysine/Laminin coated single wells in 24-well plate. NSPCs were passed when reaching about 80% confluence up to a maximum of 17-18 passages. NSPCs were usually frozen between passages 5 to 6 and stored at -80°C for further use. Experiments were performed with adult NSPCs grown as a monolayer and maintained in proliferation medium. Quiescence was induced by replacing the proliferation medium with medium containing Neurobasal-A supplemented with B27, Glutamax, human FGF-2 (20 ng/ml) and BMP4 (50 ng/ml, 5020-BP-010, RnD System). Cells were maintained in quiescence medium for 3 days before analysis. Quiescence was reversed by plating the quiescent NSPCs in normal proliferating medium. For some experiments requiring substantial amounts of cells (metabolomics and SeaHorse analyses) NSPCs were isolated form the SVZ of the corresponding floxed (or double-floxed) mouse line, given the substantial higher amount of NSPCs that can be obtained from this region as compared to the DG (Guo et al., 2012). NSPCs were seeded (2 X 10^5^ cells/well) on 6 well plates in proliferating media pre-coated with Poly-D-lysine and experiments were conducted by transducing cultures with AAVs expressing either Cre recombinase (AAV5-eGFP-Cre, Addgene #105545) or GFP only (AAV5-eGFP, Addgene #105530). To compare proliferation rates, cells were seeded (1 x10^5^ cells/well) on glass coverslips pre-coated with Poly-D-lysine and Laminin in proliferation media supplemented with or without 5x dNTP mix, 20 mM sodium acetate or 150 µM L-Aspartic acid. After 48h, cells were treated with 20µM Edu and incubated 2h followed by fixation with 4% PFA pre-warmed at room temperature for 5 mins. Cells were washed 3 times with 1X PBS followed by EdU staining using Click-iT EdU Imaging Kit (Invitrogen).

#### Measurement of Oxygen consumption by XF96 SeaHorse microplate

Oxygen consumption rate (OCR) of intact NSPCs was determined using a Seahorse XF96 extracellular flux analyser. NSPCs were seeded in coated XF96 cell culture microplate at 22,000 cells/well in 180µL of proliferation media with 25mM Glucose. The following day, proliferation media was removed from each well and cells were washed twice with assay medium consisting of (111mM NaCl, 4.7 mM KCl, 1.25 mM CaCl2, 2mM MgSO4, 1.2 mM NaH2PO4 5mM HEPES) supplemented with 25 mM glucose. Cells were incubated in 180µl medium at 37°C in a CO2-free incubator for one hour prior to measurements. OCR was monitored upon serial applications of oligomycin (2µM), FCCP (3µM) and rotenone/antimycin A (0.5µM). The optimal FCCP concentration was determined by titration (from 0.5 to 4µM) in separate experiments. To measure OCR under palmitate feeding, cells were seeded as described above and 4 h after, the proliferation media was replaced with substrate-limited proliferating medium (5mM glucose). 45 min prior the assay, cells were washed twice with aCSF supplemented with 5mM glucose (FAO assay medium) and incubated in 150µL/well FAO assay medium in a CO2-free incubator for 30-45 min at 37°C. Just before the start of the assay the media was replaced with 170µM XF Palmitate-BSA FAO substrate.

#### Western blotting

Cells were harvested in ice cold 1X PBS and homogenized in ice-cold RIPA buffer (50 mM Tris–HCl, pH 7.4, 150 mM NaCl, 1% Triton X-100, 0.1% SDS, 0.05% sodium deoxycholate, and 1 mM EDTA including protease inhibitor cocktail mix added freshly) by triturating 20-25 times with 200µL pipette tip. The homogenized lysate was incubate at 4°C for 30 min followed by centrifugation at 10,000 rpm at 4°C for 10 min and supernatant was collected. 50µg of total protein was loaded in SDS–PAGE for separation, followed by transfer to nitrocellulose membranes, and immunoblotting using the following primary antibodies (see Key resource table for details): mouse anti-PRELID1, rabbit anti-YME1L, mouse anti-OPA1, rabbit anti-TIMM17A, rabbit anti-TOMM20 and mouse anti-SDHA.

#### Proteomics sample preparation

Prior to protein extraction cells were washed using ice cold 1X PBS. Cells were then lysed using 80 µL of 8 M Urea and stored at -80°C for further use. Protein samples were sonicated to degrade the chromatin followed by centrifugation at 20,000 g for 15 min at 4°C and the supernatant was collected. Protein concentration was measured using the Direct Detect spectrometer from Merck following the Manufacturer’s instructions and 50µg protein per sample was used for further processing. Samples were mixed with 100mM Dithiothreitol (DTT) to get the final concentration of DTT 5mM followed by incubation at 37°C for 1 h at 600rpm in a thermo mixer (Eppendorf). Samples were alkylated with 40mM Chloroacetamide (CAA) and incubated for 30 min at room temperature in the dark. Samples were then mixed with endoproteinase Lys-C (1:75 (w/w) ratio of proteinase to protein) and incubated at 37°C for 4 h. Urea concentration was diluted from 8M to 2M by adding of 50mM TEAB. Samples were incubated with trypsin (1:75 (w/w) ratio of trypsin to protein) overnight at 37°C. Samples were collected and acidified to a final concentration of 1% TFA followed by StageTip extraction. SDB-RP Stage tips were pre-wetted with 30µL 100% MeOH and cleaned with 0.1% TFA, 80% ACN before equilibration with 0.1% TFA. The peptide containing samples were loaded onto SDB-RP StageTip columns and washed once with 30µL 0.1% TFA and twice with 0.1% TFA, 80% ACN followed by drying of StageTips completely with a syringe and stored at 4°C. Prior to measurement StageTips were eluted with 30 µl 1% ammonium hydroxide in 60% ACN, dried in a vacuum concentrator and resuspended in 10 µl 5% FA in 2% ACN. Samples were analysed on a Q-Exactive Plus (Thermo Scientific) mass spectrometer that was coupled to an EASY nLC 1000 UPLC (Thermo Scientific). 3 µl resuspended peptides were loaded onto an in-house packed analytical column (50 cm × 75 μm I.D., filled with 2.7 μm Poroshell EC120 C18, Agilent) and equilibrated in solvent A (0.1% FA). Peptides were chromatographically separated at a constant flow rate of 250 nL/min using the following gradient: 5-30% solvent B (0.1% formic acid in 80% acetonitrile) within 65 min, 30-50% solvent B within 13 min, followed by washing and column equilibration. The mass spectrometer was operated in data-dependent acquisition mode. The MS1 survey scan was acquired from 300-1750 m/z at a resolution of 70,000. The top 10 most abundant peptides were isolated within a 2 Da window and subjected to HCD fragmentation at a normalized collision energy of 27%. The AGC target was set to 5e5 charges, allowing a maximum injection time of 110 ms. Product ions were detected in the Orbitrap at a resolution of 17,500. Precursors were dynamically excluded for 20 s. Mass spectrometry and bioinformatic data analysis were performed by the CECAD Proteomics facility.

#### Bioinformatic MS data analysis

All mass spectrometric raw data were processed with Maxquant (version 1.5.3.8) using default parameters (Tyanova et al., 2016). Briefly, MS2 spectra were searched against the Uniprot MOUSE.fasta database, including a list of common contaminants. False discovery rates on protein and PSM level were estimated by the target-decoy approach to 0.01% (Protein FDR) and 0.01% (PSM FDR), respectively. The minimal peptide length was set to 7 amino acids and carbamidomethyolation at cysteine residues was considered as a fixed modification. Oxidation (M) and Acetyl (Protein N-term) were included as variable modifications. The match-between runs option was enabled. LFQ quantification was enabled using default settings. The Maxquant output was processed as follows: Protein groups flagged as „reverse“, „potential contaminant“ or „only identified by site“ were removed from the proteinGroups.txt. LFQ values were log2 transformed. Proteins with less than 3 valid values in at least one group of 4 replicates were removed. Missing values were replaced by imputation from a normal distribution (width 0.3, down shift 1.8). Sample t-test was used to determine significantly changing protein levels (q-value <0.05, S0 = 0.2) and a permutation-based FDR was calculated to correct for multiple testing. Enrichment of Gene Ontology, KEGG and GSEA was assessed using 1D annotation enrichment. Moreover, the obtained data was uploaded into the Ingenuity Pathway Analysis (IPA) software (Qiagen) to identify canonical pathways and gene networks that are significantly changed.

#### C13-incorporation measurements

For U-^13^C_6_ glucose (Cambridge Isotope Laboratories, CLM-1396) and U-^13^C_16_ palmitate (Cambridge Isotope Laboratories, CLM-409) chase, NSPCs were seeded in T25 flask in proliferating media containing normal (^12^C) Glucose (25mM). After 48 h unlabelled proliferating media was replaced with 25 mM U-^13^C_6_-labelled glucose or 170 µM U-^13^C_16_-labelled Palmitate media and incubated for 30 min. Cells were washed twice and harvested in ice cold saline. Cell were spin down at 2500 rpm for 5 min at 4°C and pellets were stored at -80°C until further use.

#### Whole-cell total metabolite analysis

NSPCs were seeded in T25 flask pre-coated with Poly-D-lysine and Laminin in proliferation media (supplemented with 5x dNTP mix, 20 mM sodium acetate or 150 µM L-Aspartic acid when indicated). After 48 h cells were washed twice and harvested in ice cold saline. Cell were spin down at 2500 rpm for 5 min at 4°C and pellets were stored at -80°C until further use.

#### Metabolite extraction for Liquid Chromatography mass spectrometry (LC-MS)

Metabolite extraction from each cell pellet was performed using 1 mL of a mixture of 40:40:20 [v:v:v] of pre-chilled (-20°C) acetonitrile:methanol:water (Optima^TM^ LC/MS grade, Fisher Scientific). The samples were subsequently vortexed until the cell pellets were fully suspended, before incubating them on an orbital mixer at 4°C for 30 min at 1500 rpm. For further disintegration, samples were sonicated for 10 min in an ice cooled bath-type sonicator (VWR, Germany) before centrifuging them for 10 min at 21100x g and 4°C. The metabolite-containing supernatant was collected in fresh tubes and concentrated to dryness in a Speed Vac concentrator (Eppendorf). The protein-containing pellets were collected and used for protein quantification (BCA Protein Assay Kit, Thermo Fisher Scientific).

#### LC-MS analysis of isotope-enrichments in amino acids after ^13^C-glucose or ^13^C palmitate feeding

For amino acid analysis a benzoylchlorid derivatization method (Wong et al., 2016) was used. In brief: The dried metabolite pellets was re-suspended in 20 µl LC-MS-grade waters (Optima^TM^ LC/MS grade, Fisher Scientific). The re-suspended sample were vortexed and 10 µl of 100 mM sodium carbonate (Sigma), followed by 10 µl 2% benzoylchloride (Sigma) in acetonitrile (Optima-Grade, Fisher-Scientific) were added. Samples were well vortexed before centrifuging them for 10 min 21.300x g at 20°C. Clear supernatants were transferred to fresh auto sampler tubes with conical glass inserts (Chromatographie Zubehoer Trott) and analyzed using an Acquity iClass UPLC (Waters) connected to a Q-Exactive HF (Thermo) mass spectrometer (MS). For the analysis of the amino acids 1 µL of the derivatized samples was injected onto a 100 x 1.0 mm HSS T3 UPLC column (Waters). The flow rate was set to 100 µL/min using a buffer system consisted of buffer A (10 mM ammonium formate (Sigma), 0.15% formic acid (Sigma) in LC-MS-grade water (Optima^TM^ LC/MS grade, Fisher Scientific) and buffer B (acetonitrile, Optima-grade, Fisher-Scientific). The LC gradient was: 0% B at 0 min; 0-15% B 0-0.1 min; 15-17% B 0.1-0.5 min; 17-55% B 0.5-14 min, 55-70% B 14-14.5 min; 70-100% B 14.5-18 min; 100% B 18-19 min; 100-0% B 19-19.1 min, 19.1-28 min 0% B. The mass spectrometer was operating in positive ionization mode monitoring a m/z range between 50 and 750. The heated electrospray ionization (ESI) source settings of the mass spectrometer were: Spray voltage 3.5kV, capillary temperature 250°C, sheath gas flow 60 AU and aux gas flow 20 AU at a temperature of 250°C. The S-lens was set to a value of 60 AU. Data analysis of isotope ratios was performed using the TraceFinder software (Version 4.2, Thermo Fisher Scientific). Identity of each compound was validated by authentic reference compounds, which were analysed independently. For the isotope enrichment analysis the area of the extracted ion chromatogram (XIC) of each isotope [M + H]^+^ was determined with a mass accuracy (<5 ppm) and a retention time (RT) precision (<0.1 min), before calculating the proportions of each detected isotope towards the sum of all isotopes of the corresponding compound. These proportions were given as percent values for each isotope.

#### GC-MS analysis of isotope-enrichments in metabolites from glycolysis and TCA cycle after ^13^C-glucose or ^13^C palmitate feeding

Similar to the analysis of the isotope enrichment analysis in the amino acids, isotope enrichment analysis of glycolysis and TCA cycle metabolites were determined using gas chromatography (GC) coupled to a high resolution, accurate mass MS (Q-Exactive GC-Orbitrap, Thermo Fisher Scientific). For this analysis metabolites were derivatized using a two-step procedure starting with an methoxyamination (methoxyamine hydrochlorid, Sigma) followed by a trimethyl-silylation using N-Methyl-N-trimethylsilyl-trifluoracetamid (MSTFA, Macherey-Nagel). Dried samples were re-suspended in 5 µL of a freshly prepared (20 mg/mL) solution of methoxyamine in pyridine (Sigma) to perform the methoxyamination. These samples were then incubated for 90 min at 40°C on an orbital shaker (VWR) at 1500 rpm. In the second step additional 45 µL of MSTFA were added and the samples were incubated for additional 30 min at 40°C and 1500 rpm. At the end of the derivatisation the samples were centrifuged for 10 min at 21.100x g and 40 µL of the clear supernatant were transferred to fresh auto sampler vials with conical glass inserts (Chromatographie Zubehoer Trott). For the GC-MS analysis 1 µL of each sample was injected using a PAL autosampler system (Thermo Fisher Scientifc) using a Split/Splitless (SSL) injector at 300 °C in splitless-mode. The carrier gas flow (helium) was set to 2 ml/min using a 30m DB-35MS capillary column (0.250 mm diameter and 0.25 µm film thickness, Agilent). The GC temperature program was: 2 min at 85°C, followed by a 15°C per min ramp to 330°C. At the end of the gradient the temperature was held for additional 6 min at 330°C. The transfer line and source temperature were both set to 280°C. The filament, which was operating at 70 eV, was switched on 2 min after the sample was injected. During the whole gradient period the MS was operated in full scan mode covering a m/z range between 70 and 800 with a scan speed of 20 Hertz. For data analysis peak areas of XICs of each isotope of a compound-specific fragment [M - e^-^]^+^ were determined using the TraceFinder software (Version 4.2, Thermo Fisher Scientific). XICs were extracted with a mass accuracy (<5 ppm) and a RT precision (<0.05 min), as compared to independently analysed authentic reference compounds. Subsequently proportions of each detected isotope towards the sum of all isotopes of the corresponding compound-specific fragment were determined. These proportions are given as percent values for each isotope. The flux into each compound was analysed from a single compound-specific fragment. Accordingly, 3-phosphoglyceric acid was analysed from a three carbon-containing fragment with the chemical formula C14H36O7PSi4 and m/z 459.12702. Phosphenolpyruvic acid was analysed from a three carbon-containing fragment with the chemical formula C11H26O6PSi3 and m/z 369.07693. Pyruvic acid was analysed from a three carbon-containing fragment with the chemical formula C6H12NO3Si and m/z 174.05809. Lactic acid was analysed from a three carbon-containing fragment with the chemical formula C8H19O3Si2 and m/z 219.08672. Citric acid was analysed from a five carbon-containing fragment with the chemical formula C11H21O4Si2 and m/z 273.09729. Alpha-ketoglutaric acid was analysed from a five carbon-containing fragment with the chemical formula C8H12NO3Si and m/z 198.058096. Succinic acid was analysed from a four carbon-containing fragment with the chemical formula C9H19O4Si2 and m/z 247.08164. Fumaric acid was analysed from a four carbon-containing fragment with the chemical formula C9H17O4Si2 and m/z of 247.08164. Malic acid was analysed from a four carbon-containing fragment with the chemical formula C9H17O4Si2 and m/z of 247.08164.

#### Anion-Exchange Chromatography Mass Spectrometry (AEX-MS) of the analysis of Nucleotides

For the analysis and quantification of xy, the dried pellet of the metabolite extract was re-suspended in 100 µL of H_2_O (LC-MS Optima-Grade, Thermo Scientific) and analysed using a Dionex ionchromatogrypy system (ICS 5000, Thermo Scientific). The applied protocol was adopted from (Schwaiger et al., 2017). In brief: 10 µL of polar metabolite extract were injected in full loop mode using an overfill factor of 3, onto a Dionex IonPac AS11-HC column (2 mm × 250 mm, 4 μm particle size, Thermo Scientific) equipped with a Dionex IonPac AG11-HC guard column (2 mm × 50 mm, 4 μm, Thermo Scientific). The column temperature was held at 30°C, while the auto sampler was set tot 6°C. A potassium hydroxide gradient was generated by the eluent generator using a potassium hydroxide cartridge that was supplied with deionized water. The metabolite separation was carried at a flow rate of 380 µL/min, applying the following gradient. 0-3 min, 10 mM KOH; 3-12 min, 10−50 mM KOH; 12-19 min, 50-100 mM KOH, 19-21 min, 100 mM KOH, 21-22 min, 100-10 mM KOH. The column was re-equilibrated at 10 mM for 8 min. The eluting metabolites were detected in negative ion mode using ESI MRM (multi reaction monitoring) on a Xevo TQ (Waters) triple quadrupole mass spectrometer applying the following settings: capillary voltage 2.75 kV, desolvation temp. 550°C, desolvation gas flow 800 L/h, collision cell gas flow 0.15 mL/min. All peaks were validated using two MRM transitions one for quantification of the compound, while the second ion was used for qualification of the identity of the compound. Data analysis and peak integrationand quantification was performed using the TargetLynx Software (Waters).

#### NAD/NADH measurement

NSPCs (2 x 10^6^) were harvested in cold PBS. Cells were spin down at 2000 rpm for 5 min at 4 °C and cell pellets were collected for NAD and NADH measurements by using an NAD/NADH colorimetric Assay Kit (Abcam) according to the manufacturer’s instructions.

#### RNA sequencing

NSPCs were harvested in TRIzol reagent (Invitrogen) and RNA was isolated according to the manufacturer’s instructions. Libraries were prepared using the Illumina® Stranded TruSeq® RNA sample preparation Kit. Library preparation started with 2µg total RNA. ERCC RNA Spike-In Mix that provides a set of external RNA controls was added to the total RNA prior to library preparation to enable performance assessment. After poly-A selection (using poly-T oligo-attached magnetic beads), mRNA was purified and fragmented using divalent cations under elevated temperature. The RNA fragments underwent reverse transcription using random primers. This was followed by second strand cDNA synthesis with DNA Polymerase I and RNase H. After end repair and A-tailing, indexing adapters were ligated. The products were then purified and amplified (14 PCR cycles) to create the final cDNA libraries. After library validation and quantification (Tape Station 4200), equimolar amounts of library were pooled. The pool was quantified by using the Peqlab KAPA Library Quantification Kit and the Applied Biosystems 7900HT Sequence Detection System. The pool was sequenced Illumina NovaSeq6000 sequencing instrument with a PE100 protocol.

#### Bioinformatic RNA-Seq data analysis

Quality control, trimming, and alignment or raw data were performed using the “nf-core” RNA-seq pipeline (v3.0) (Ewels et al., 2020). The reference genome sequence and transcript annotation used were Mus Musculus GRCm39 ensembl v103. Differential expression analysis was performed in R (v4.0.3) (https://www.R-project.org/) with “DESeq2” package (v1.30.1) (Love et al., 2014) and Log (Fold Change) shrinkage estimation was calculated with “apeglm” (v1.12.0) (Zhu et al., 2019). Only genes with a minimum coverage of 10 reads in 6 or more samples from each pairwise comparison were considered as candidates for differential expression analysis. Genes were considered deferentially expressed if they showed a |*log*2(*Fold Change*)| > 0.5 and if they were below the p-value threshold. Nominally significant p-values were considered with a p-value < 0.05 without multiple testing correction. Expression level data for heatmaps are in length Scaled TPM (Transcripts Per Kilobase Million) units.

## Quantification and statistical analysis

Data are represented as means ± SE. Graphical illustrations and significance were obtained with GraphPad Prism 7 (GraphPad). Significance was calculated as described in each figure legend.

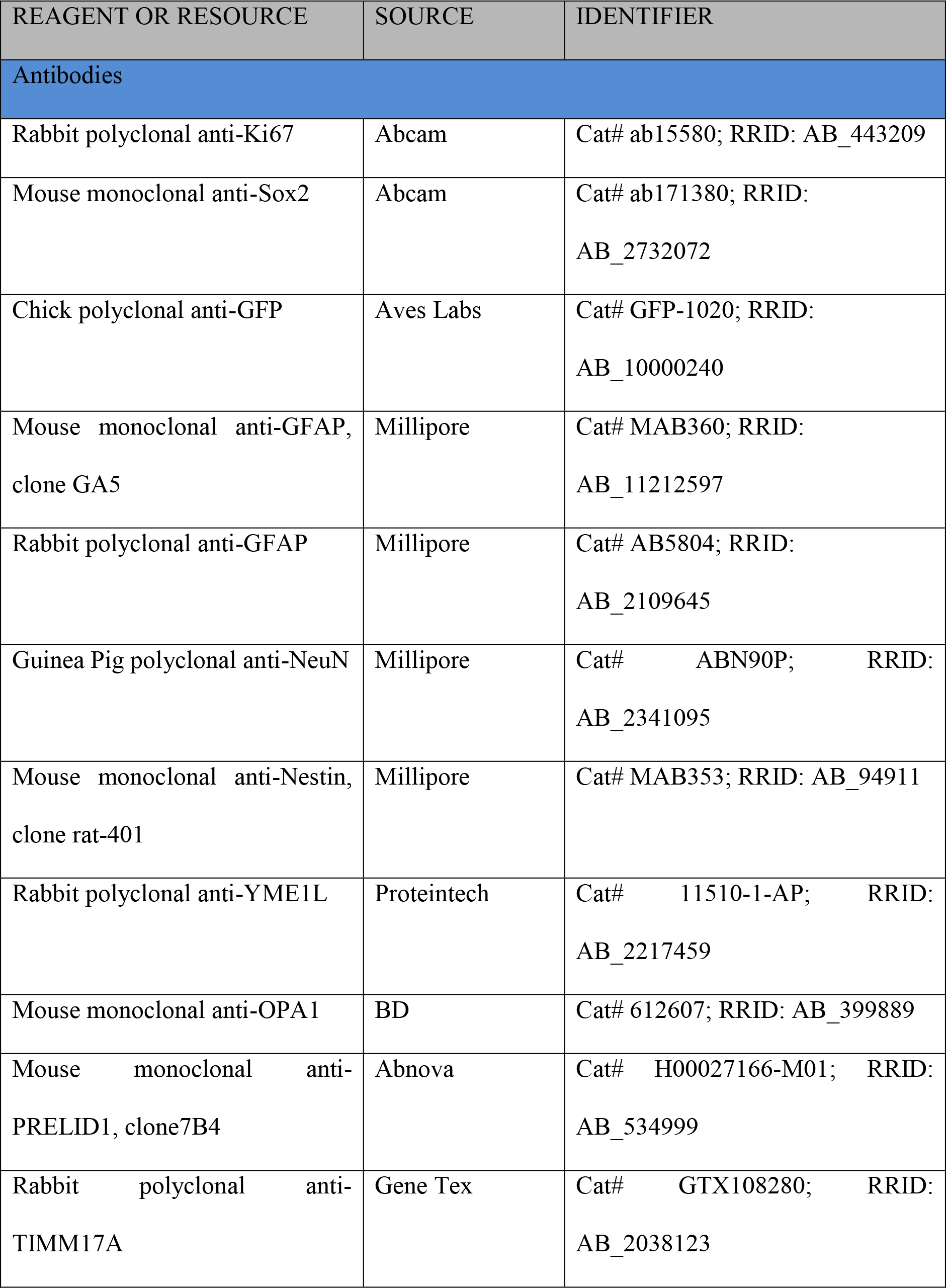
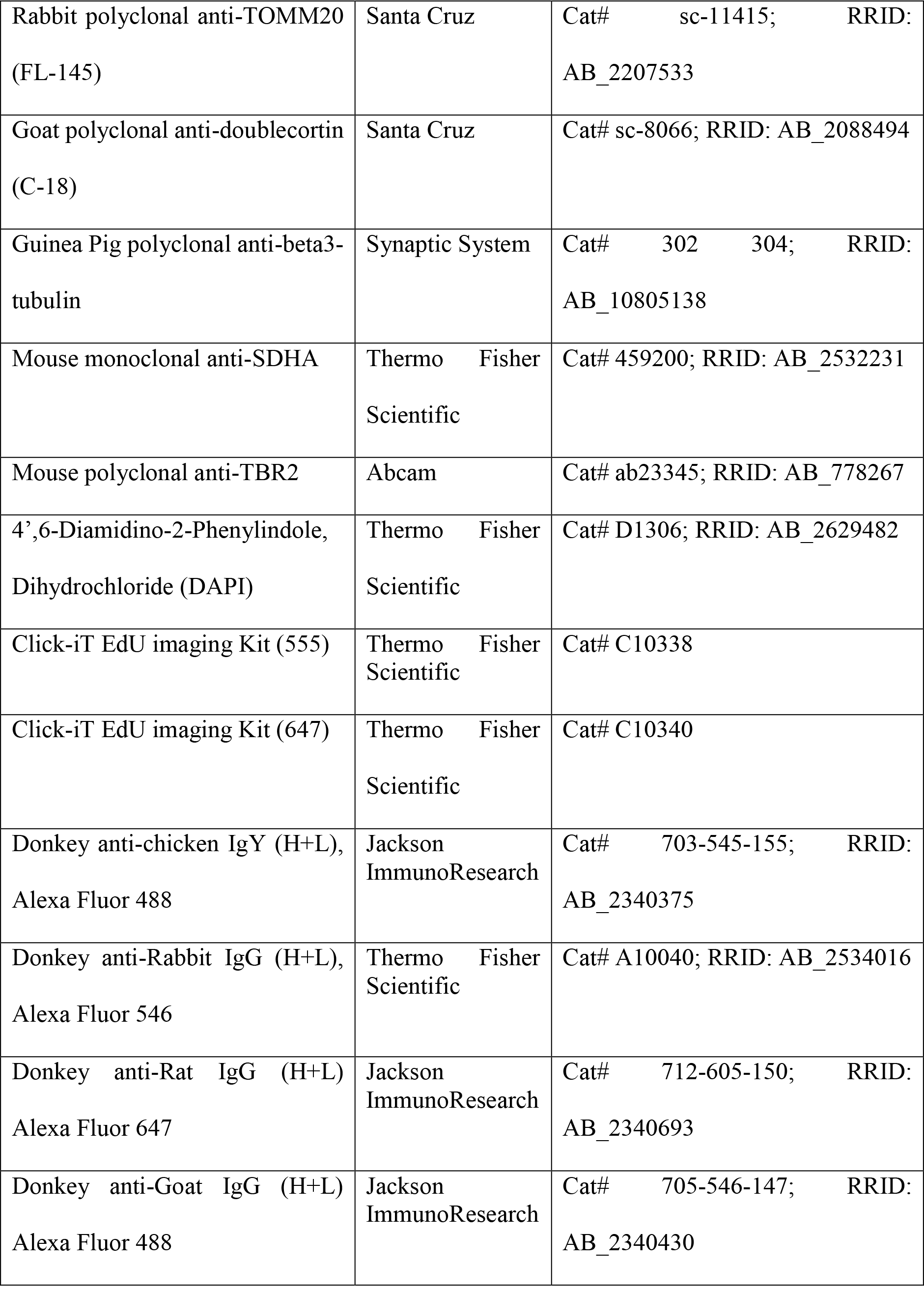
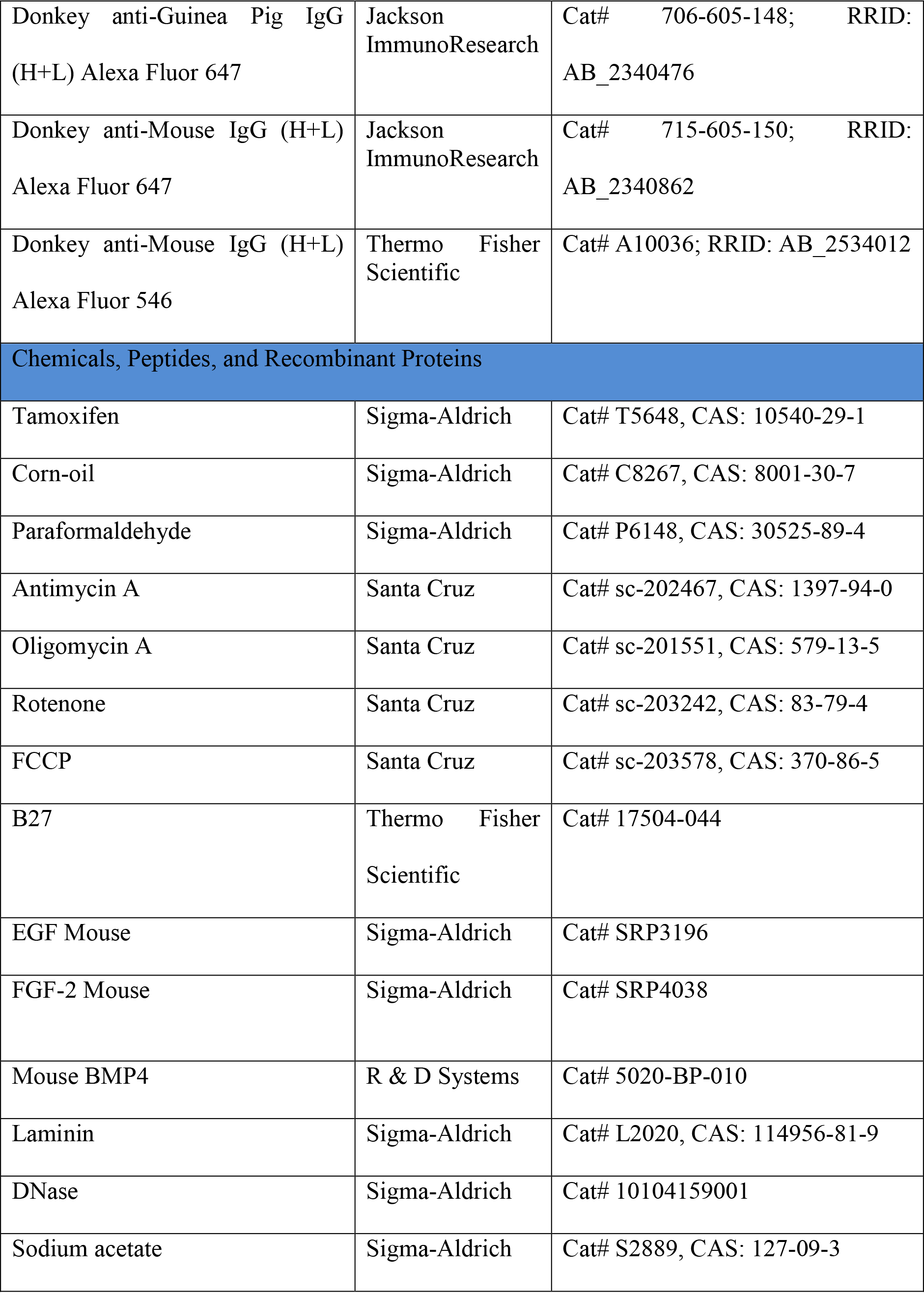
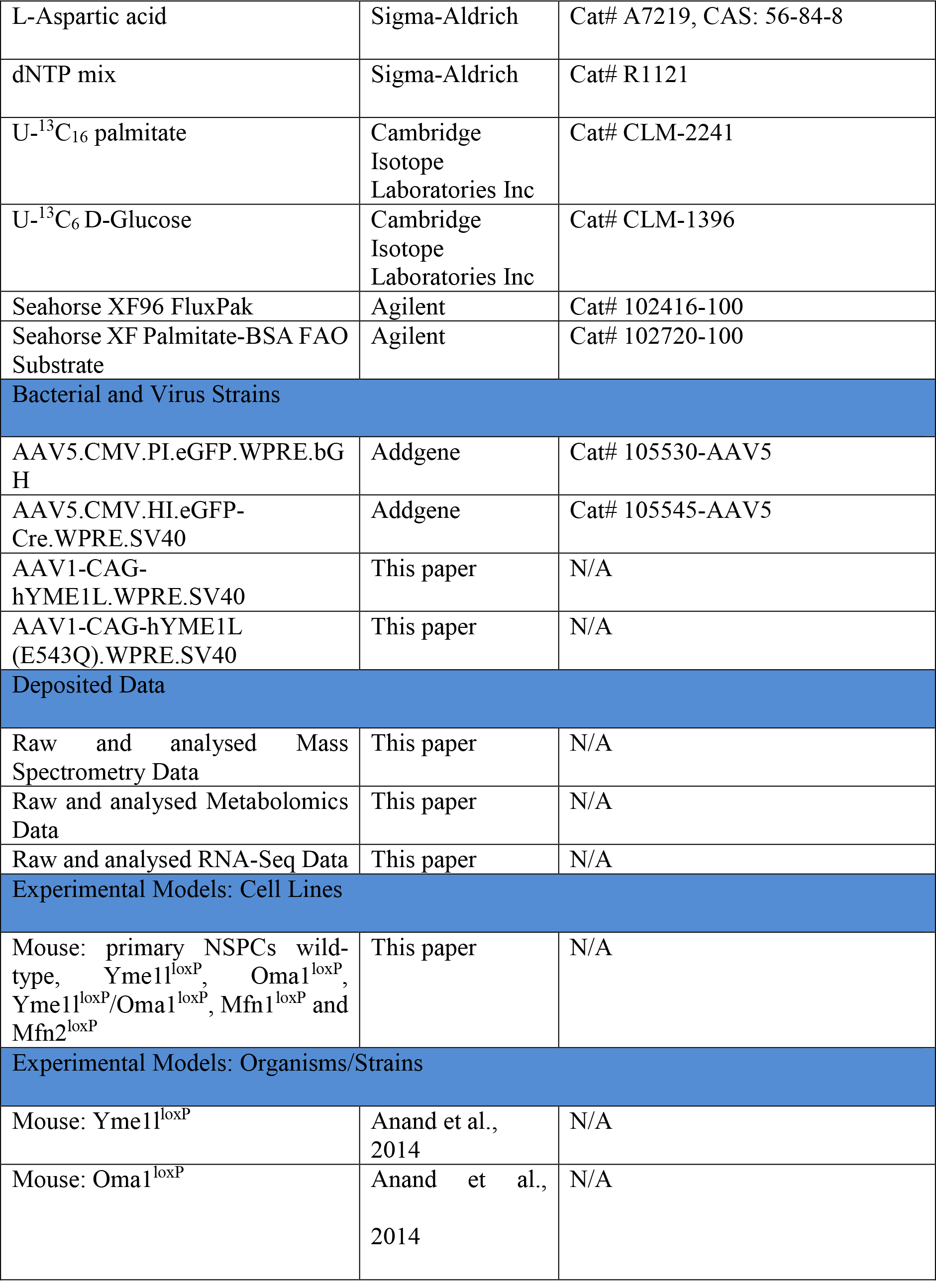
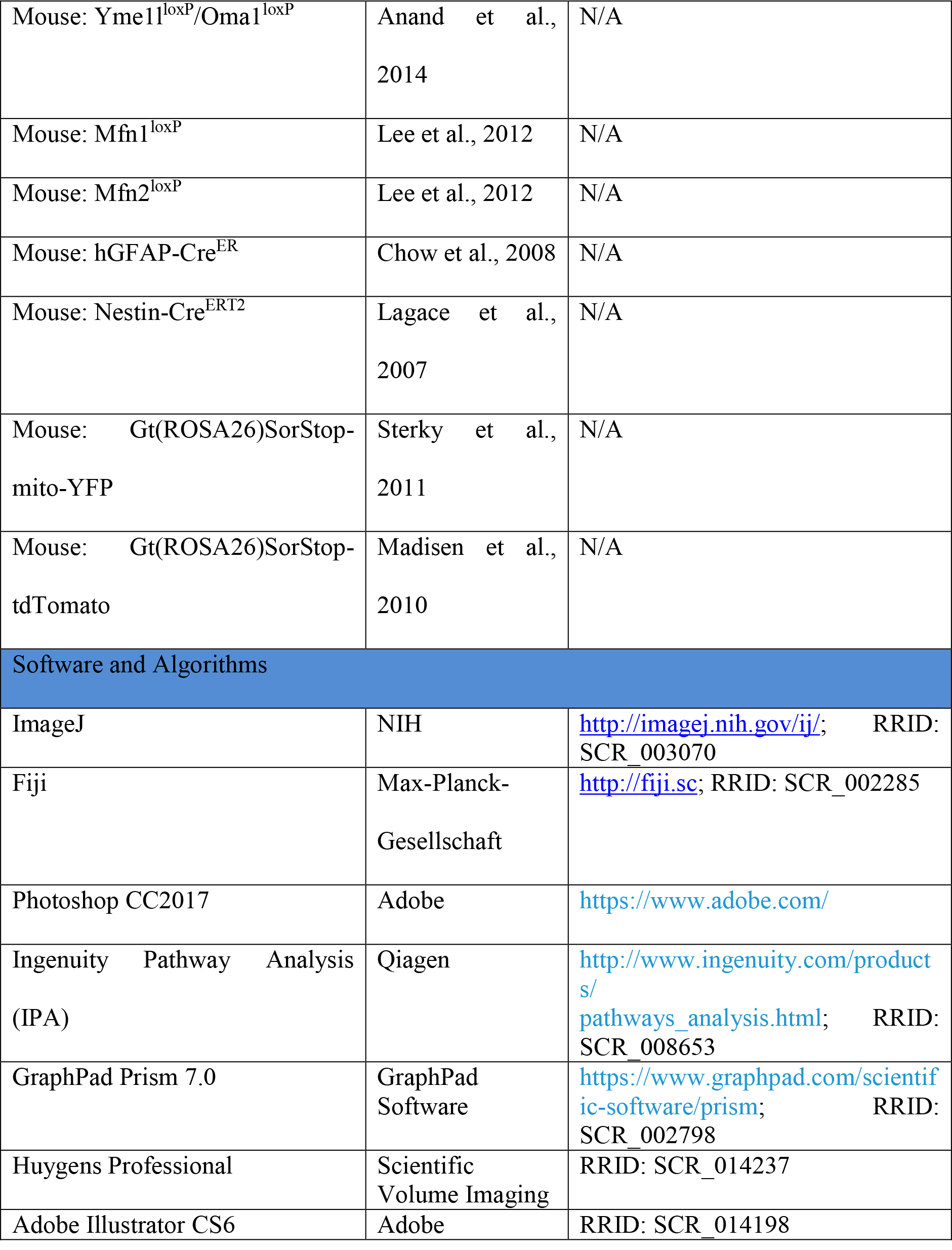

## Legend to Supplemental Figures

**Figure S1. Related to Figures 1 and 2.**
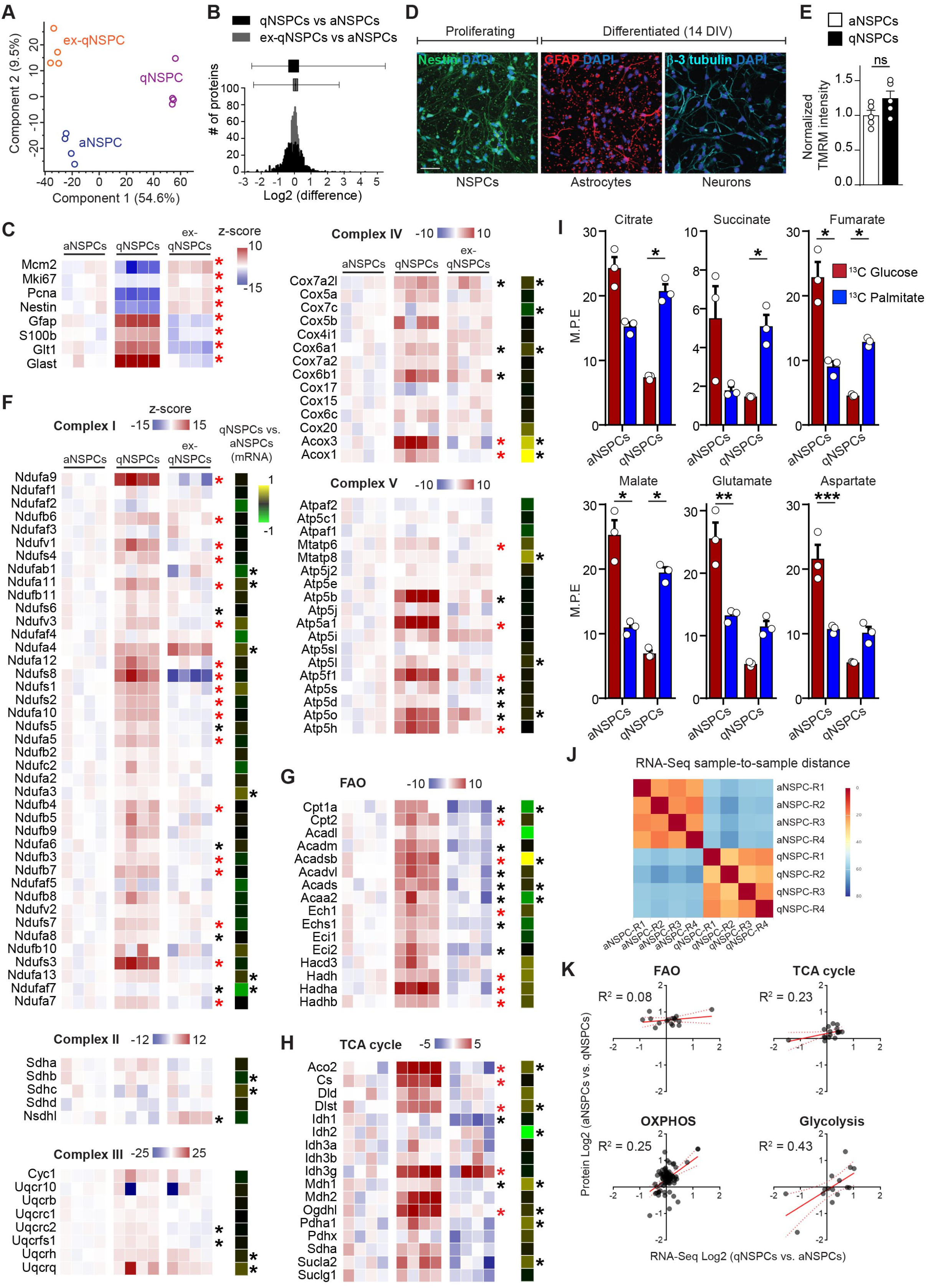
Assessment of NSPC state-dependent mitochondrial changes in proteome and metabolic fluxes. **(A)** PCA plot of aNSPCs, qNSPCs and ex-qNSPCs proteomic datasets. **(B)** Graph depicting the proteome-wide changes in protein expression levels of qNSPCs (in black) and ex-qNSPCs (in gray) with respect to aNSPCs, illustrating a wider distribution in the case of qNSPCs. **(C)** Heat maps of normalized LFQ intensities showing the differential expression of selected markers linked to proliferation (Mcm2, Mki67, Pcna, Nestin) as well as quiescence (Gfap, S100b, Slc1a2, Slc1a3) in aNSPCs and qNSPCs. Significant changes (n= 4 independent experiments per condition; FDR-adjusted ≤ 0.05) between qNSPCs and both aNSPCs and ex-NSPCs are indicated with a red asterisk. **(D)** Example of NSPC cultures isolated from the adult hippocampal DG and maintained either in active proliferation (Nestin+, left panel) or exposed to differentiation media for 14 days (following gradual withdrawal of grow factors), in which NSPCs acquire either stellate astrocytic (GFAP+) morphologies or a neuronal identity (Beta-3 tubulin+). Bar, 50 µm. **(E)** Quantification of mitochondrial membrane potential (following incubation with 25 μM TMRM) in aNSPCs and qNSPCs (n= 5 experiments; Welch’s t-test). **(F)** Heat-maps of normalized LFQ intensities showing the differential expression (z-score) of electron transport chain (OXPHOS) subunits forcomplexes I-V in aNSPCs, qNSPCs and ex-qNSPCs. Significant changes (n= 4 independent experiments per condition; FDR-adjusted ≤ 0.05) are indicated with an asterisk (red asterisk: qNSPCs significant vs both aNSPCs and ex-NSPCs; black asterisk: qNSPCs significant vs only one other category). The right column reports on the fold change of mRNA levels of the corresponding genes between qNSPCs and aNSPCs (n= 4 independent experiments per condition; FDR-adjusted ≤ 0.05). **(G-H)** Heat-maps of normalized LFQ intensities showing the differential protein expression of FAO (G) and TCA cycle (H) enzymes in aNSPCs, qNSPCs and ex-qNSPCs. Significant changes (n= 4 independent experiments per condition; FDR-adjusted ≤ 0.05) are indicated with an asterisk (red asterisk: qNSPCs significant vs both aNSPCs and ex-NSPCs; black asterisk: qNSPCs significant vs only one other category). The right column reports on the fold change of mRNA levels of the corresponding genes between qNSPCs and aNSPCs (n= 4 independent experiments per condition; FDR-adjusted ≤ 0.05). **(I)** Graphs showing the mass isotopomer enrichment analysis for the indicated TCA cycle metabolites and amino acids in aNSPCs and qNSPCs (n= 3 independent experiments per condition; Holm-Sidak’s t-test) after either ^13^C6-Glucose (in red) or ^13^C6-Palmitate (in blue) supplementation. M.P.E., Molar Percent Enrichment (calculated for each isotope). **(J)** Clustered heat-map showing the sample-to-sample distance relative to the RNA-Seq data obtained from aNSPCs and qNSPCs (n= 4 independent experiments per condition; FDR-adjusted ≤ 0.05). **(K)** Plots illustrating the extent of correlation (R^2^) between protein and mRNA levels of the genes associated to FAO, TCA cycle, OXPHOS (Complexes subunits shown in E) and glycolysis (data obtained from n= 4 independent experiments per condition and dataset). Means ± SEM; *, *P* < 0.05; **, *P* < 0.01; ***, *P* < 0.005.

**Figure S2. Related to Figures 1, 2 and 3.**
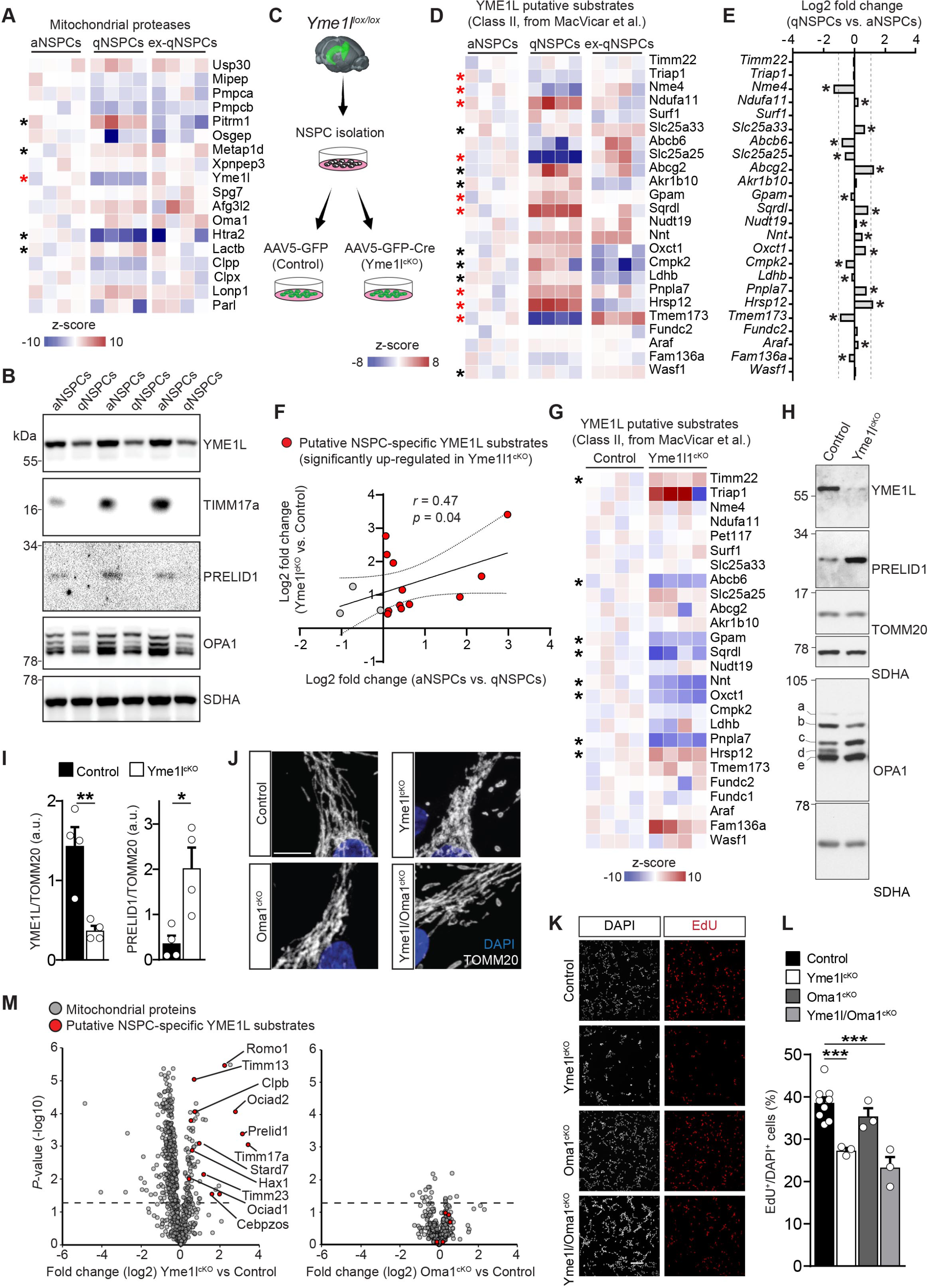
Assessment of YME1L substrate specificity via Yme1l^cKO^ NSPCs. **(A)** Heat-maps of normalized LFQ intensities of all detected mitochondrial proteases in NSPCs. Significant changes (n= 4 independent experiments; FDR-adjusted ≤ 0.05) are indicated with an asterisk (red asterisk: qNSPCs significant vs both aNSPCs and ex-NSPCs; black asterisk: qNSPCs significant vs only one other category). **(B)** Immunoblot of YME1L as well as of two of its validated proteolytic targets (TIMM17a and PRELID1) in qNSPCs and aNSPCs. OPA1 expression levels and SDHA loading controls are shown. **(C)** Experimental design illustrating the conditional deletion of *Yme1l* in adult NSPCs maintained *in vitro* via treatment with GFP-Cre or GFP-only AAVs. **(D)** Heat-maps of normalized LFQ intensities showing the differential expression of class II putative YME1L substrates. Significant changes (n= 4 independent experiments; FDR-adjusted ≤ 0.05) are indicated with an asterisk (red asterisk: qNSPCs significant vs both aNSPCs and ex-NSPCs; black asterisk: qNSPCs significant vs only one other category). **(E)** Fold change of mRNA levels of class II putative YME1L substrates between qNSPCs and aNSPCs (n= 4 independent experiments; FDR-adjusted ≤ 0.05). **(F)** Scatter plot showing class I putative YME1L substrates that underwent a significant accumulation in Yme1l^cKO^ NSPCs. Substrates represented by a red dot indicate those that were also found at reduced protein levels in qNSPCs as compared to aNSPCs. **(G)** Heat maps of normalized LFQ intensities showing the differential expression of class II putative YME1L substrates in Yme1l^cKO^ NSPCs. Asterisks indicate significant changes (n= 4 independent experiments; FDR-adjusted ≤ 0.05). **(H)** Immunoblots of NSPCs few days after treatment with the GFP-Cre AAV (Yme1l^cKO^) showing the ablation of YME1L as well as the accumulation of the YME1L substrate PRELID1 and the increased OMA1-mediated OPA1 processing leading to the expected accumulation of S-OPA1 isoforms c and e. **(I)** Quantification of YME1L and PRELID1 expression levels in Yme1l^cKO^ NSPCs as shown in B (n= 4 independent experiments, Unpaired t-test). **(J)** Confocal pictures showing the morphology of mitochondria (immunolabeled against Tomm20) in control, Yme1l^cKO^, Oma1^cKO^ and Yme1l/Oma1^cKO^ NSPCs. Bar, 10 µm. **(K)** Examples of NSPCs belonging to the indicated genotypes and showing the extent of cell proliferation following 2h treatment with the base analog EdU (20µM). **(L)** Graph reporting on the quantification of NSPC proliferation as shown in E (n= 3-9 independent experiments, One-way Anova followed by Holm-Sidak’s correction). **(M)** Volcano plots showing the changes in mitochondrial proteomes of Yme1l^cKO^ and Oma1^cKO^ NSPCs compared to controls (n= 4 independent experiments per genotype). NSPC-specific class I YME1L substrates are highlighted in red. Cut-off line set at –log10 (*P*-value) = 1.3. Means ± SEM; *, *P* < 0.05; **, *P* < 0.01; ***, *P* < 0.005.

**Figure S3. Related to Figures 3 and 4.**
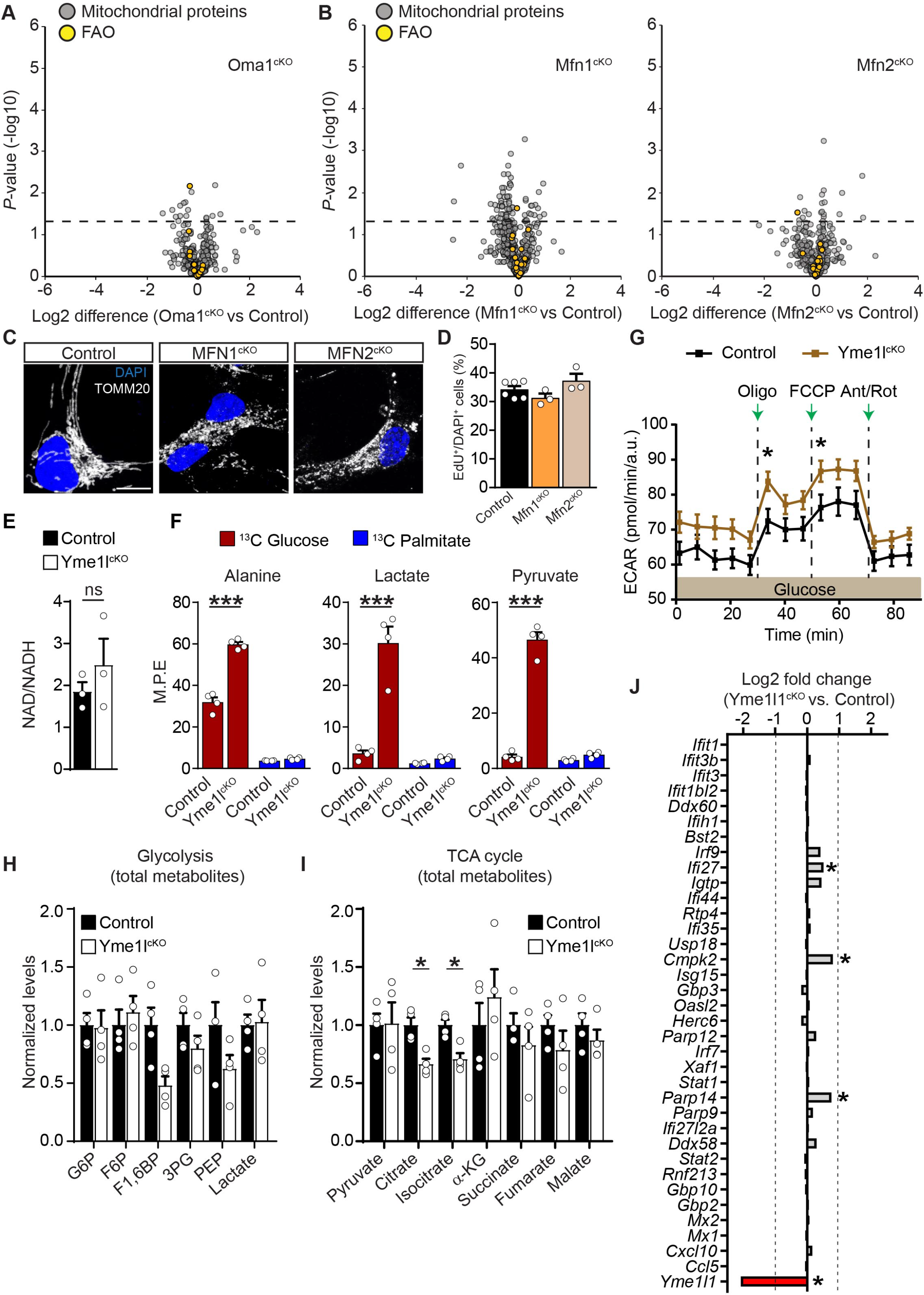
Metabolic analysis of Yme1l^cKO^ NSPCs. **(A)** Volcano plot showing the changes in the mitochondrial proteome of Oma1^cKO^ NSPCs compared to controls (n= 4 independent experiments per genotype). FAO enzymes are highlighted in yellow. Cut-off line set at –log10 (*P*-value) = 1.3. **(B)** Volcano plots showing the changes in the mitochondrial proteome of Mfn1^cKO^ and Mfn2^cKO^ NSPCs compared to controls (n= 3 independent experiments per genotype). FAO enzymes are highlighted in yellow. Cut-off line set at –log10 (*P*-value) = 1.3. **(C)** Representative pictures of NSPCs obtained from control, Mfn1^cKO^ and Mfn2^cKO^ immunostained for the mitochondrial marker TOMM20. Bar, 10 µm. **(D)** Proliferation assay in control, Mfn1^cKO^ and Mfn2^cKO^ NSPCs (n= 3-6 independent experiments per genotype). **(E)** Measurement of NAD/NADH ratio in control and Yme1l^cKO^ NSPCs (n= 3 independent experiments). **(F)** Graphs showing the mass isotopomer enrichment analysis for the indicated metabolites in control and Yme1l^cKO^ NSPCs (n= 4 independent experiments per condition; Holm-Sidak’s t-test) after either ^13^C6-Glucose (in red) or ^13^C6-Palmitate (in blue) supplementation. M.P.E., Molar Percent Enrichment (calculated for each isotope). **(G)** Extracellular acidification rate (ECAR) measurement in control and Yme1l^cKO^ NSPCs fed with glucose (n= 30 repetitions, 3 independent experiments; Holm-Sidak multiple t-test). **(H)** Measurement of glycolytic metabolites in Yme1l^cKO^ NSPCs, normalized to control NSPCs (4 independent experiments; Holm-Sidak multiple t-test). **(I)** Measurement of TCA cycle metabolites in Yme1l^cKO^ NSPCs, normalized to control NSPCs (4 independent experiments; Holm-Sidak multiple t-test). **(J)** Transcriptomic analysis in NSPCs of interferon-stimulated genes previously described for being induced in Yme1l^cKO^ MEFs (4 independent experiments; FDR-adjusted ≤ 0.05). Means ± SEM; *, *P* < 0.05; **, *P* < 0.01; ***, *P* < 0.005.

**Figure S4. Related to Figure 5.**
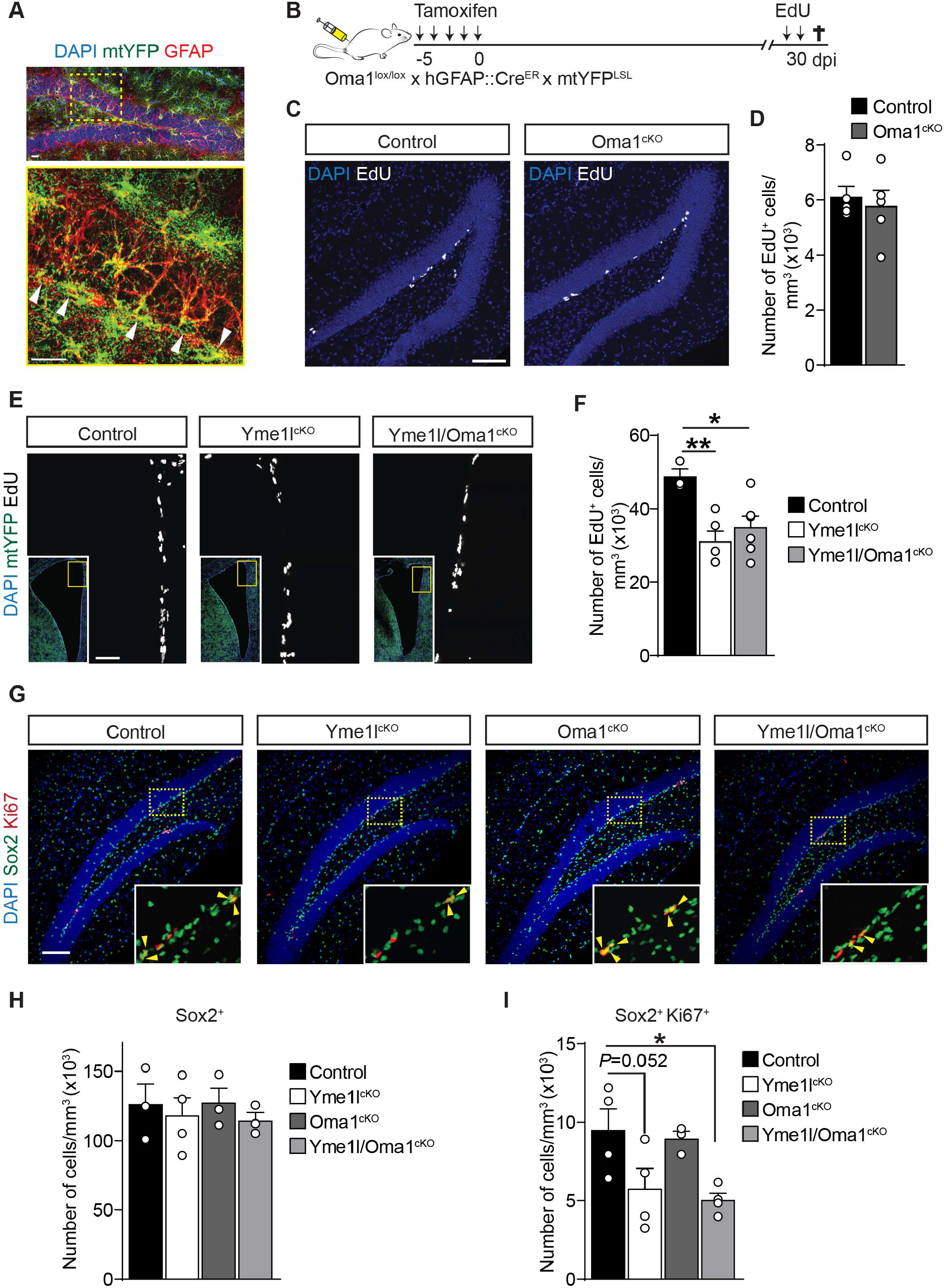
Lack of YME1L impairs NSPC proliferation in the adult DG and SVZ. **(A)** Section of the DG of an hGFAP::Cre^ER^ x mtYFP^LSL^ mouse after tamoxifen administration, showing the extent of recombination (on the basis of the mtYFP reporter gene expression) in GFAP+ cells lining the SGZ (arrowheads). Bar, 20 µm. **(B)** Experimental design illustrating the tamoxifen-induced conditional deletion of *Oma1* in hGFAP::Cre^ER^ x mtYFP^LSL^ mice. **(C)** Examples EdU labelling in the DG of the indicated genotypes. Bar, 70 µm. **(D)** Quantification of NSPC proliferation for the indicated genotypes (n= 5 mice per group; Unpaired t-test). **(E)** Examples EdU labelling in the SVZ lining the lateral ventricle of the indicated genotypes. Insets show an overview of the lateral ventricle. Bar, 50 µm. **(F)** Quantification of NSPC proliferation in the SVZ for the indicated genotypes (n= 3-6 mice per group; One-way Anova followed by Holm-Sidak’s multiple comparison test). **(G)** Examples showing the DG of the indicated genotypes and the density of Sox2^+^ cells (identifying both radial glia-like NSPCs and astrocytes) as well as that of Ki67^+^ (proliferating) cells. Insets show enlargements of the boxed areas. Arrowheads point to double positive (Sox2^+^/ki67^+^) cells. Bar, 100 µm. **(H)** Quantification of Sox2^+^ cells (including both radial glia-like NSPCs and astrocytes) in the SGZ of the DG of the indicated genotypes (n= 3-4 mice per group, One-way Anova followed by Holm-Sidak’s multiple comparison test). **(I)** Quantification of dividing Sox2^+^/ki67^+^ cells in the SGZ of the DG of the indicated genotypes (n= 3-4 mice per group, One-way Anova followed by Holm-Sidak’s multiple comparison test). Means ± SEM; *, *P* < 0.05; **, *P* < 0.01.

**Figure S5. Related to Figure 5.**
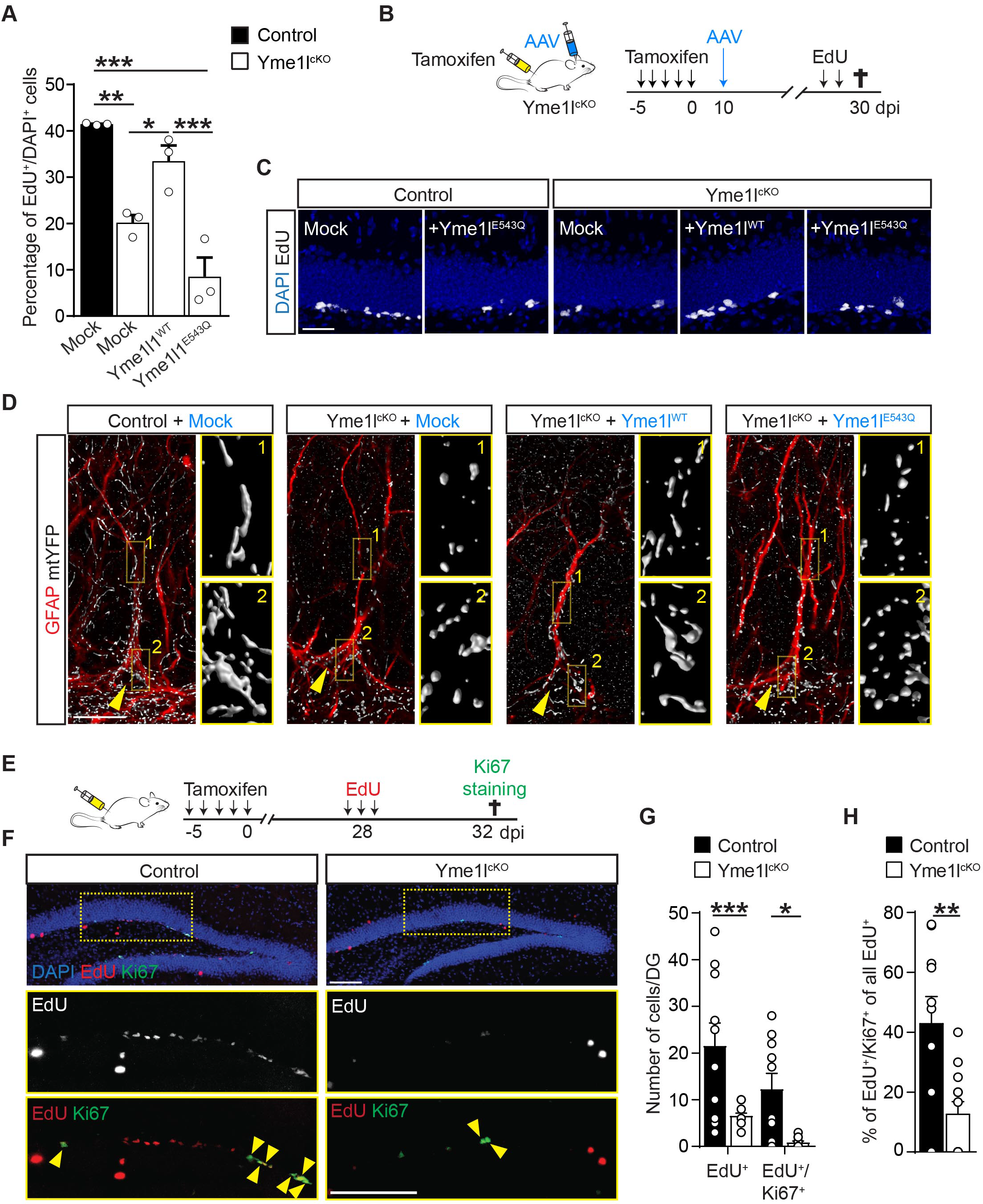
Regulation of NSPC proliferation by YME1L is independent from changes in mitochondrial dynamics. **(A)** Quantification of control or Yme1l^cKO^ NSPCs proliferation *in vitro* following transduction with the indicated AAVs and treatment with EdU (n= 3 independent experiments; one-way Anova followed by Holm-Sidak’s multiple comparison test). **(B)** Experimental design illustrating the tamoxifen-induced conditional deletion of *Yme1l* in hGFAP::Cre^ER^ x mtYFP^LSL^ mice followed by stereotactic delivery of AAVs encoding for Yme1l1^WT^ or the proteolytically inactive variant Yme1l1^E543Q^. EdU treatments were performed briefly before sacrifice to assess changes in NSPC proliferation. **(C)** Examples of EdU-treated animals for the indicated genotypes and conditions to assess NSPC proliferation in the DG. Bar, 40 µm. **(D)** Examples of individual radial glia-like mtYFP^+^/GFAP^+^ NSPCs in the DG of the indicated genotypes and following injection of mock, Yme1l1^WT^ or Yme1l1^E543Q^-encoding AAVs, showing the corresponding changes in mitochondrial morphology. Arrowheads points to the cell soma. Right panels show zooms of the boxed areas in the cell soma and along the main radial process. Bar, 15 µm. **(E)** Experimental design illustrating the tamoxifen-induced conditional deletion of *Yme1l* in hGFAP::Cre^ER^ x mtYFP^LSL^ mice followed by EdU treatments (which incorporates into dividing cells) 4 days before sacrifice and assessment of still-dividing EdU+ cells by immunostaining against the proliferation marker Ki67. **(F)** Examples of EdU-treated animals as depicted in E for the indicated genotypes showing proliferating EdU^+^/Ki67^+^ cells in the DG. Low panels show enlargements of the boxed areas and report on individual and merged channels. Arrowheads point to double positive (EdU^+^/Ki67^+^) cells in the SGZ. Bar, 100 µm. **(G)** Quantification of total EdU^+^ and EdU^+^/Ki67^+^ cells in the SGZ of control and Yme1l^cKO^ mice (n= 10-12 mice, two-way Anova). **(H)** Proportion of EdU^+^/Ki67^+^ cells of all EdU^+^ cells (n= 10-12; unpaired t-test). Means ± SEM; *, *P* < 0.05; **, *P* < 0.01; ***, *P* < 0.005.

**Figure S6. Related to Figure 6.**
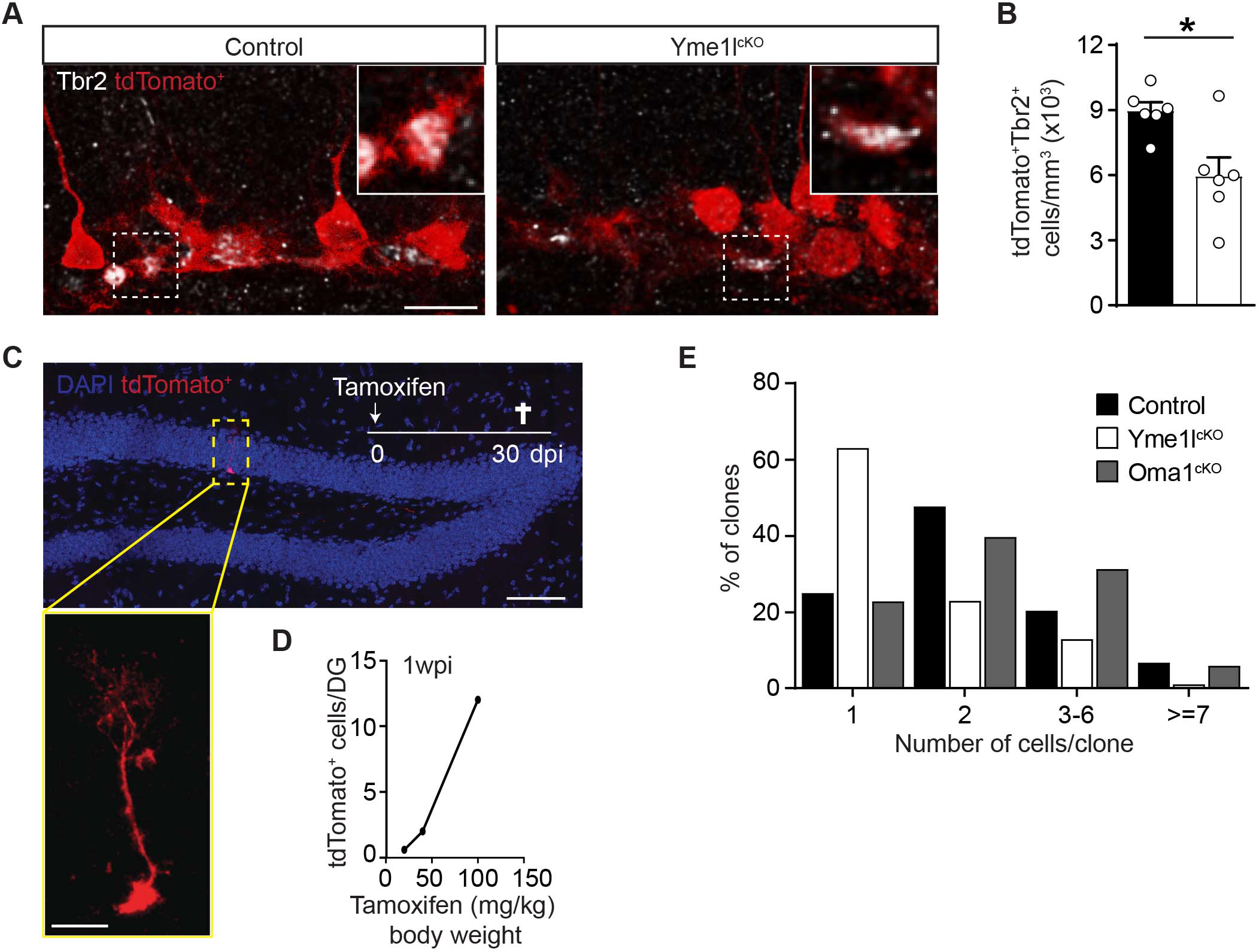
Analysis of type II NSPCs and clonal analysis in Yme1l^cKO^ NSPC. **(A)** Examples of Tbr2 immunostaining in the DG of Yme1l^lox/lox^ x Nestin::Cre^ERT2^ x tdTomato^LSL^ and control mice. Insets report zooms of the boxed areas. Bar, 20 µm. **(B)** Quantification of TdTomato^+^ and Tbr2^+^ cells in Yme1l^cKO^ mice (n= 6 mice per group, unpaired t-test). **(C)** Example of a single recombined TdTomato+ radial glia-like NSPC following a single low dose tamoxifen administration. The low panel shows an enlargement of the boxed area. Bars, 80 and 25 µm. **(D)** Tamoxifen dose-dependent increase in the number of TdTomato^+^ cells found in the whole DG of Nestin::Cre^ERT2^ x TdTomato^LSL^ mice by 1 week after induction. **(E)** Quantification of clone size per genotype (n= 7-9 mice per genotype). Means ± SEM; *, *P* < 0.05.

## Notes

### Competing Interest Statement

The authors have declared no competing interest.

